# An Engineering Explanation of Human Vertical Reaching: Optimal Trajectory Planning of Asymmetric Velocity Profiles in the Vertical Plane via a Softplus-Based Generalized Dynamic Jerk Cost Function

**DOI:** 10.64898/2026.10.03.756410

**Authors:** Muhamad Braa Kabbani, Alaa Marouf

## Abstract

The results in this article demonstrate that the central nervous system (CNS) plans trajectories in the vertical plane by minimizing a Generalized Dynamic Jerk (GJerk) cost function, **which we have proposed as an advancement of the traditional Minimum Jerk cost function**. While the traditional Jerk cost function generates smooth trajectories by minimizing the third derivative of joint angle over time, it neglects real dynamic constraints—most notably the effect of gravity. According to our engineering interpretation, the arm functions as a constrained manipulator where gravity acts as a non-linear, state-dependent bias; thus, the asymmetry in velocity profiles is not an anomaly but an optimal adaptation to the gravitational field. Consequently, it predicts mathematically symmetrical upward and downward movements, even in contexts where gravitational forces play a decisive role. The proposed GJerk cost function extends the classical framework by incorporating position-dependent dynamic coefficients into the cost function, thereby embedding gravitational effects directly within the planning process. As a result, the generated movements are not only smooth in a mathematical sense but also physically and mechanically consistent with the actual torque and energy demands imposed by the musculoskeletal system. The GJerk cost function reduces angular change in high-gravitational-torque regions because it makes the joint configuration with the highest gravitational torque more expensive within the cost function, prompting the planner to choose a slower and more conservative movement. This does not physically dissipate energy, but it acts as a behavioral damping mechanism, similar to how humans naturally slow down when approaching a high-load or potentially dangerous configuration. In this way, the GJerk cost function reduces abrupt changes in joint kinetic energy and prevents sudden jumps in acceleration, ultimately producing a smoother and more stable trajectory that aligns with both biomechanical reality and observed human behavior. Within this framework, upward movements become inherently more costly than downward ones, penalizing configurations that require large torques to counteract gravity. This induces a biomechanically meaningful asymmetry, ensuring the resulting trajectories reflect the body’s true mechanical constraints rather than mere geometric smoothness. Our empirical results demonstrate that the GJerk cost function provides a more realistic and physiologically grounded account of motor planning in the vertical plane, linking smoothness optimization with genuine dynamic requirements and offering a closer approximation to biological motor behavior

## Introduction: Motor Planning, Optimal Control Theory, and Gravity Integration in Upper Limb Kinematics

The study of human movement planning and the neural and biomechanical strategies used to guide the upper limb is a cornerstone of research in biomechanics and Optimal Control Theory (OCT). This theoretical framework provides the mathematical lens through which we understand how the brain selects a single, efficient trajectory from an infinite set of possibilities—a problem known as the ‘degrees of freedom’ redundancy. Methodological foundations in this field were established in the early 1980s by Soechting and Lacquaniti [1], [2]. They introduced the concept of ‘invariant characteristics,’ identifying motor features that persist regardless of changes in movement speed, load, or amplitude. During this foundational era, research focused heavily on the horizontal plane, where the influence of gravity is neutralized by the plane of motion. Observational studies by Morasso (1981) [24] and Abend et al. (1982) [25] demonstrated a strict kinematic regularity characterized by straight hand paths and temporally symmetric, “bell-shaped” velocity profiles. These descriptive observations led to the formulation of the first formal control laws, such as the Minimum Hand Jerk cost function [10]. This cost function assumes that the central nervous system (CNS) prioritizes absolute smoothness by minimizing the third derivative of position, successfully predicting the temporal symmetry observed in gravity-free horizontal reaches [11], [15], [17].

However, the transition from planning in the horizontal plane to the vertical plane reveals a deeper mechanical and neural complexity that disrupts the traditional symmetry of the bell-shaped velocity profile. While gravity is perpendicular to movement in the horizontal plane—thereby preserving its classic symmetric bell-shaped form—Atkeson and Hollerbach (1985) [3] proved that in the sagittal plane, gravity becomes a primary determinant of the trajectory, inevitably deforming the profile into an asymmetrical bell-shaped velocity profile. Subsequent observational research by Papaxanthis et al. (2003) [4], [28] identified clear ‘Directional Asymmetries’; specifically, vertical movements exhibit a distinct velocity asymmetry, where they lack the temporal symmetry found in horizontal reaches. In upward movements (against gravity), the CNS tends to significantly shorten the acceleration duration compared to the deceleration phase. This deliberate manipulation, as explained by Papaxanthis et al. [28] and Gaveau and Papaxanthis (2011) [31], aims to exploit gravity as a natural braking force, further accentuating this velocity asymmetry. By shortening acceleration, the brain generates sufficient momentum to overcome gravity, subsequently allowing gravity to “assist” in deceleration, thereby reducing the need for active antagonist muscle activation and preserving the energetic efficiency of the limb [27].

Since 2010, scientific research has demonstrated that this asymmetric velocity profile is not merely a passive physical response; rather, it is the result of a ‘neural integration’ of an internal gravity model embedded within the brain [21], [23]. A pivotal contribution to this field was established by Berret et al. (2011) through a series of foundational studies [14], [16]. In their initial work [14], they proposed the ‘Inactivation Principle’—a dynamic control law explaining how the Central Nervous System (CNS) selects trajectories by minimizing of Energy.

Subsequently, by applying Inverse Optimal Control (IOC) in their follow-up research [16], the authors provided definitive evidence that human motor planning is governed by a ‘composite cost function.’ This framework demonstrates that the brain does not optimize for a single objective; instead, it reconciles competing constraints by integrating the minimization of angle jerk, to ensure kinematic smoothness, with the minimization of Energy, to maximize energetic efficiency. The complexity of this integration lies in the CNS’s ability to weight these constraints dynamically. By balancing the mathematical penalty assigned to angle jerk against the physiological cost of mechanical work, the brain effectively solves an optimization problem that accounts for both the desired path fluidity and the gravitational potential energy. This integrated cost function provides the mechanistic basis for the observed periods of ‘muscle silence’ (gravitational free-fall) during downward movements, where the CNS deliberately ceases active torque production to exploit gravity. Consequently, the work of Berret et al. stands as a cornerstone reference, confirming that the internal gravity model functions as a sophisticated computational framework. This framework not only reconciles the kinematic demand for smoothness with the physiological demand for efficiency but also characterizes the CNS as an active, predictive optimizer that treats gravity as a predictable variable rather than a stochastic disturbance in the motor planning process.

Following this foundation, the post-2011 era marked a ‘Golden Age’ of empirical expansion, largely building upon the theoretical lineage of Bastin’s school. A critical milestone was the work of Gaveau et al. (2021) [21], who provided cross-species evidence that the neural integration of gravity for motor optimization is a fundamental, conserved feature of vertebrate neural architecture. This empirical robustness was further solidified across functional domains: Gaveau et al. (2014, 2016) [22], [23] confirmed the paradigm’s sensitivity to granular gravitational torques at the shoulder, elbow, and wrist, while Poirier et al. (2022) [40] anchored the paradigm in inhibitory neurobiology. Furthermore, Russo et al. (2020, 2021) [47], [48] generalized the framework to whole-body modular synergies. The paradigm also demonstrated remarkable plasticity; robotic studies by Gaveau et al. (2016) [51], S. F. et al. (2021) [55], and Macaluso et al. [44] revealed rapid reoptimization under environmental perturbations. Its cognitive validity was confirmed by Hilt et al. [19], [43], [49] in free-endpoint reaching, and by Lebon et al. [56] and Desoche et al. [57] during mental motor imagery.

Despite the extensive empirical validation of the Bastin-Berret Paradigm, the field has reached a state of theoretical stagnation. For the past fifteen years, research has remained tethered to the 2011 formulation, treating deviations—such as the breakdown of effort-minimization under high-accuracy constraints [41] or the influence of subjective cognitive priorities [19]—as isolated anomalies rather than integrated features.

Recent findings (2021–2024), particularly by Poirier et al. [40], [41] and Bucchieri et al. [42], reveal that motor control is governed by ‘Flexible Optimality,’ where the CNS dynamically reweights accuracy, effort, and gravity in real-time. These observations are fundamentally incompatible with the static mathematical architecture of the 2011 model, leading to a literature that is empirically rich but theoretically fragmented.

To resolve this impasse, we introduce the GJERK (Generalized Dynamic Jerk) model. Unlike the classical paradigm, which relies on static, additive constants, GJERK elevates the cost function into a gravity-weighted dynamic operator. This model finally bridges the gap between 1980s mechanical regularities and the flexible, adaptive intelligence observed in modern motor control, transforming the cost function from a rigid constraint into a context-aware mechanism.

## Materials and Methods

### 1. Theoretical Foundation of the Proposed GJERK Model

In the jerk classical framework, the movement cost function for reaching movements is expressed as the time-integral of the squared third derivative of position (squared jerk):

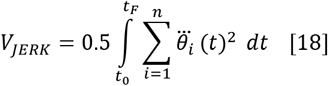

This model assumes that the Central Nervous System (CNS) seeks uniform smoothness across the kinematic chain, but it completely ignores environmental physics and gravity, making it unable to explain why humans “freeze” certain joints or exploit “free fall” during downward movements.

The GJERK model conceptualizes motor control not as the minimization of an abstract smoothness parameter, but as a dynamic integration of kinematics and gravitational dynamics. Unlike traditional frameworks that treat movement smoothness as a uniform goal, we propose that the Central Nervous System (CNS) performs a context-aware optimization where gravitational torque *G*_*i*_(*θ*), acts as a dynamic modulator of the cost functional. The core optimization objective is defined as:

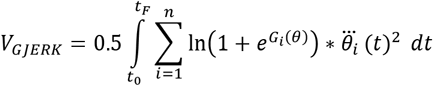

In this formulation, 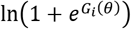 serves as a weight that adjusts the penalty of the angular jerk 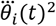 in real-time. This modulation implies that the CNS does not minimize jerk uniformly across the kinematic chain; rather, it distributes ‘jerk variance’ through a smooth, continuous weighting function based on the gravitational state of each joint, effectively transforming the cost function into a gravity-weighted operator.

The model’s predictive power emerges from the mechanical implications of the gravitational torque sign and magnitude. When a joint experiences assistive gravitational torque G < 0, the gravitational potential energy facilitates motion, thereby reducing the cost of movement. Consequently, the CNS identifies these joints as preferred actuators, allocating the majority of the kinematic variance—and thus the highest jerk—to these segments. This strategy exploits gravity to enhance energetic efficiency, explaining the fluid, gravity-led trajectories observed in downward reaches.

Conversely, in neutral configurations where *G* > 0, the system reverts to a baseline kinematic smoothness minimization, governed primarily by inertial constraints. However, the model’s most significant contribution lies in its treatment of resistive gravity where *G* > 0 represents a proportional penalty on acceleration changes leading to a balanced distribution of smoothness, whereas *G* ≫ 0 imposes a severe penalty on acceleration changes; in these conditions, the CNS adopts a strategy of differential smoothness allocation where the joint becomes mechanically filtered by its own gravitational loading, acting as a fluid anchor, providing a formal mechanistic basis for synergies such as the observed “freezing” of segments, arising not as an active neural restriction but as a consequence of optimizing within a high-potential-energy landscape. Ultimately, by employing the continuous softplus modulation of 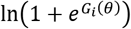 nand relying solely on the physical modulation of the gravitational torque, the GJERK model provides a unified framework that characterizes the human body as an integrated system that exploits environmental physics to achieve efficient and stable motion.

This mechanism creates distinct synergistic behaviors based on the gravitational states *G*_*i*_(*θ*)and *G*_*i*_(*θ*): when both joints are assistive *G*_*i*_(*θ*), *G*_*i*+1_(*θ*)< 0), the CNS utilizes both equally to maximize fluidity; conversely, when both are resistive (*G*_*i*_(*θ*), *G*_*i*+1_(*θ*) ≫ 0), the system adopts a “freezing” synergy where, if *G*_*i*_(*θ*)> *G*_*i*+1_(*θ*), the CNS prioritizes jerk minimization for joint *i*, effectively using it as the primary fluid anchor. In mixed configurations, the strategy is defined by the assistive joint’s role as the primary actuator: if *G*_*i*_(*θ*) < 0 and *G*_*i*+1_(*θ*) > 0, joint *i* provides the necessary motive power while the resistive joint *i* + 1, is stabilized for trajectory precision, whereas if *G*_*i*_(*θ*) > 0 and *G*_*i*+1_(*θ*) < 0, joint *i* + 1 becomes the primary actuator while the resistive joint *i* is prioritized for jerk minimization to act as a stable base. This mechanistic framework effectively explains complex biological motor patterns, such as the stabilization of the vertebral column and the rhythmic arm swinging observed during locomotion, by demonstrating how the CNS dynamically modulates joint impedance to maintain postural integrity and balance energetic costs. Ultimately, by employing the continuous softplus modulation of *G*_*i*_(*θ*), and relying solely on the physical modulation of the gravitational torque, the GJERK model provides a unified framework that characterizes the human body as an integrated system that exploits environmental physics to achieve efficient and stable motion.

### 2. Comparative Framework and Evolutionary Benchmarking

The evolutionary trajectory of cost function formulations in motor control is summarized in Table 1. This table highlights the cost function architectures originally proposed by Berret et al. [16], which have demonstrated significant predictive power and served as the benchmark for motor planning models to date. As illustrated, the literature has evolved through these distinct formulations, where the Energy expenditure term is explicitly proposed to model the Effort required by the neuromuscular system to overcome mechanical resistance.

**Table 1:**
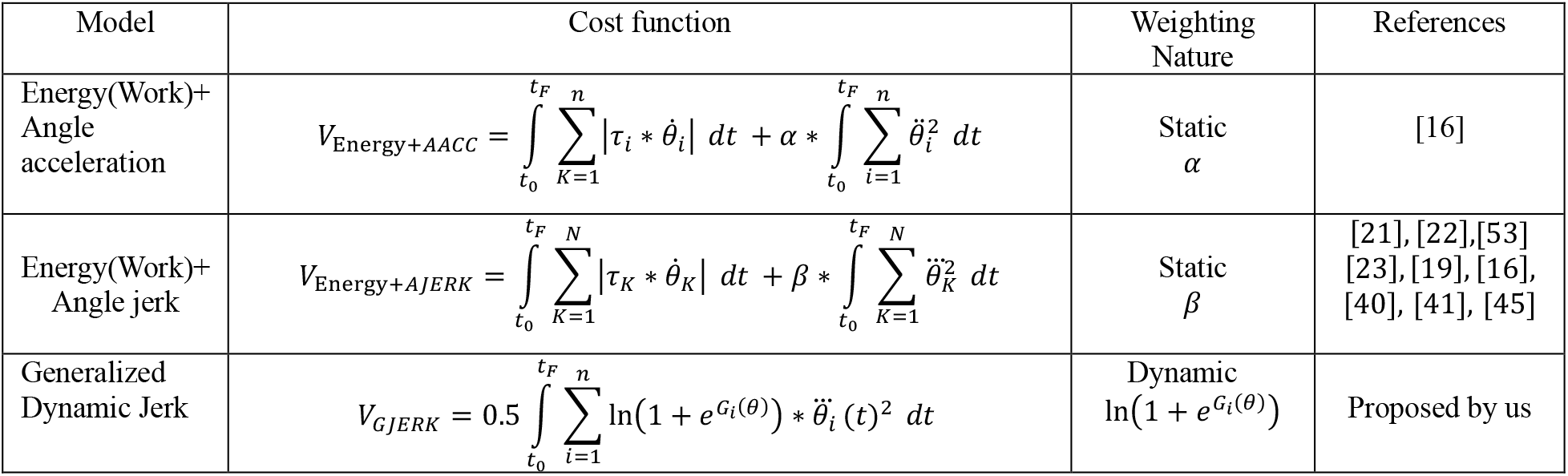
Evolution of cost function architectures in motor control modeling: From parameter-dependent composites to the GJERK gravity-weighted operator.

The fundamental distinction between these frameworks lies in their handling of environmental constraints. While Berret’s composite models rely on static weighting coefficients—requiring the CNS to perform exogenous tuning of competing objectives— the GJERK model internalizes these constraints. By substituting static weights with the continuous softplus-modulated gravitational torque 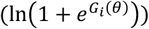, we transform the cost functional into a context-aware operator. This shift signifies that observed postural synergies and velocity asymmetries are not the result of multiple competing objectives, but rather the emergent physical consequence of a singular, gravity-weighted smoothness minimization. Consequently, GJERK provides a more parsimonious and physically grounded account of motor planning, aligning computational theory with the inherent physics of human movement.

In physics and mechanical engineering, ‘energy’ and ‘mechanical work’ are equivalent; specifically, angular mechanical work—the integral of torque over angular velocity—mathematically quantifies the energy cost of movement in biomechanics. Therefore, these terms are used interchangeably in this study to represent the cumulative mechanical effort within the objective function, as follow:

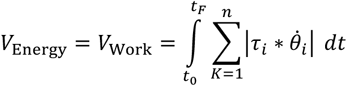

### 3. Experimental Setup and Data Acquisition

To rigorously validate the proposed GJERK framework, we conducted a comparative analysis using the empirical dataset originally established by Berret et al. [16]. This pointing task serves as an ideal benchmark for our model, as it effectively isolates the complex interaction between movement smoothness and gravitational dynamics by removing any restrictive precision constraints along the vertical axis. Unlike traditional point-to-point movements where high spatial accuracy demands minimize the discernibility between competing cost models, reaching toward a vertical bar creates a broad path tolerance. By allowing the vertical target height to remain unconstrained *Y*(*tf*)=*free*, the experimental design permits the Central Nervous System (CNS) to freely select reaching endpoints that optimize effort, thereby revealing the underlying physical and cognitive strategies for gravity-related motor planning; this conceptual divergence in behavioral outcomes is illustrated in **Figure 1**.

**Figure 1:**
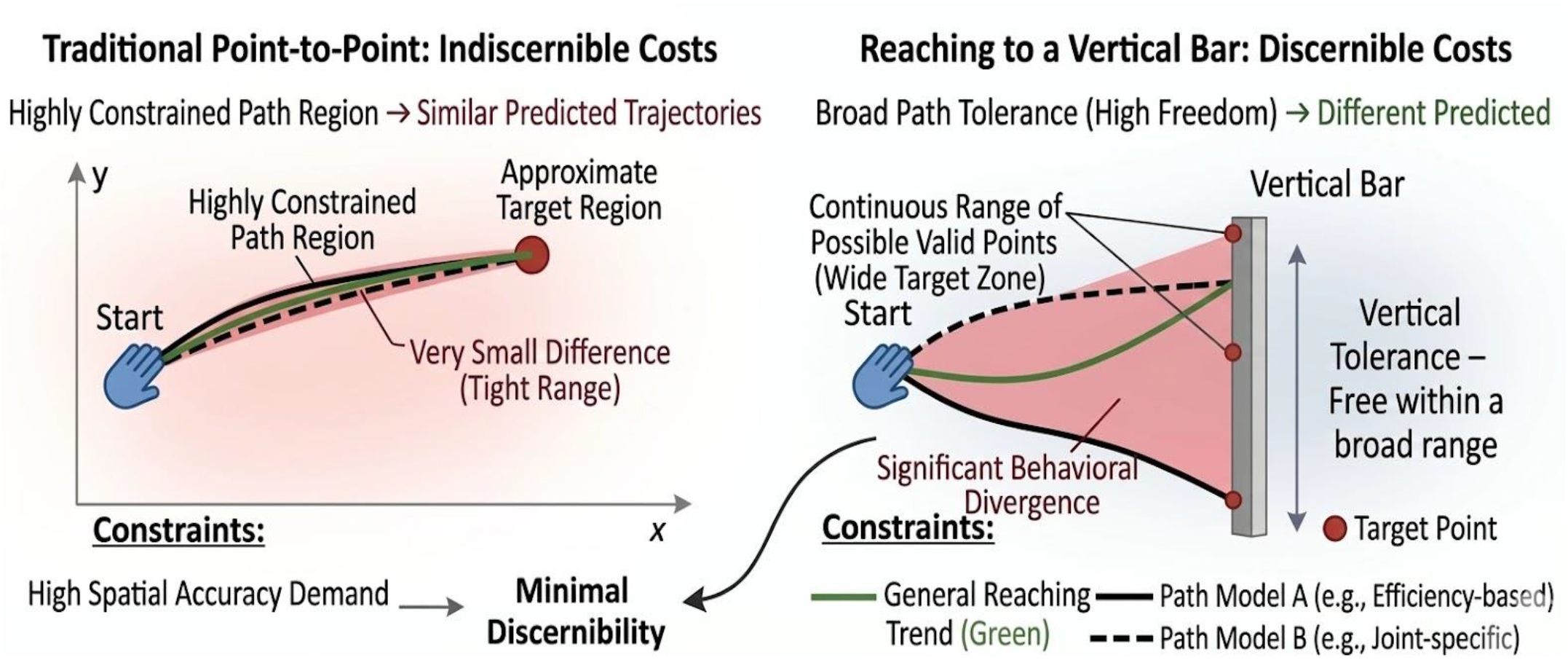
Increased Behavioral Divergence in Bar-Reaching Due to Higher Spatial Freedom.

Participants were seated comfortably and instructed to perform reaching movements toward a vertical target bar, which consisted of a 2.5-meter rigid, non-labeled tube placed in the parasagittal plane and strictly aligned with the participant’s right shoulder. To maintain consistency across participants, the horizontal distance between the shoulder and the target bar was precisely set to 85% of each participant’s total arm length (*L*_*Total*_). During the execution of the task, movement was restricted to the shoulder and elbow joints only; to ensure this constraint, the wrist joint was immobilized using lightweight, rigid splints firmly attached to both the forearm and the hand. This restriction was critical to simplify the kinematic chain to a two-degree-of-freedom system, allowing for a focused analysis of the shoulder-elbow coordination without the confounding variables introduced by wrist flexibility:

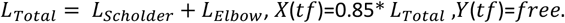

To provide a comprehensive evaluation of the model under varying mechanical conditions, five distinct initial postures (P1–P5) were systematically defined. As illustrated in **Figure 2**, these initial configurations were determined by physical reference points situated in a vertical plane approximately 10 cm lateral to the participant’s right shoulder. Each posture represented a specific state in the configuration space, defined by the following angular pairs (shoulder; elbow) measured in degrees:

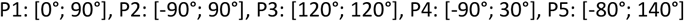

**Figure 2:**
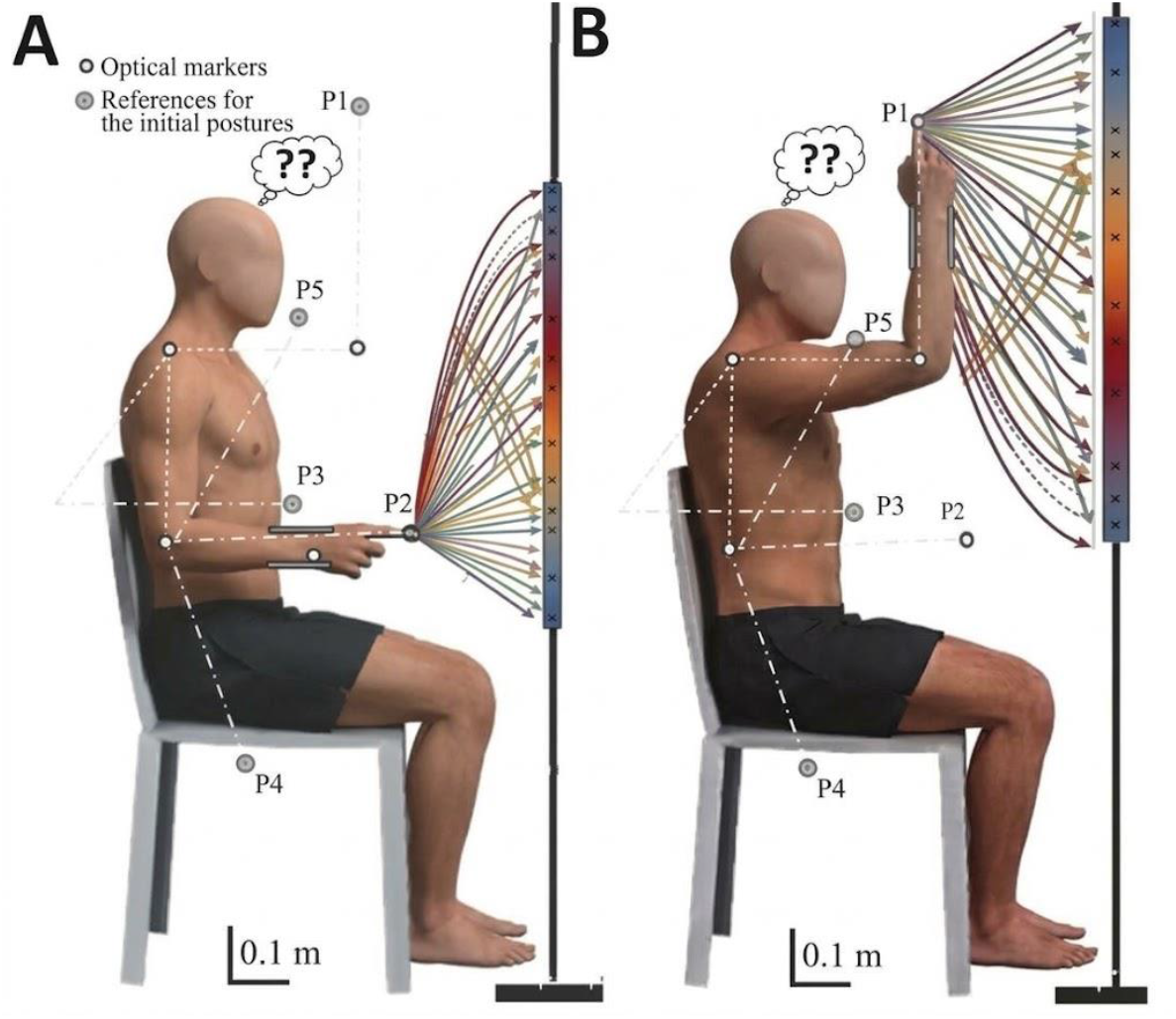
The figure illustrates how the human arm has more degrees of freedom than necessary to reach a target. In both examples, a person can choose different paths to move their hand from the starting position to the target on the bar. The question marks highlight the difficulty in predicting the precise path the central nervous system will choose.

The strategic selection of these five postures was intended to map the entire parasagittal workspace and to systematically vary the mechanical demands—specifically the gravitational torque—at the onset of each movement. By initiating reaching movements from these diverse points, the experimental setup challenges the motor system to resolve distinct inertia-gravity combinations for a uniform task goal. Consequently, this provides a rigorous test for the GJERK model, as it must demonstrate predictive accuracy across a broad spectrum of initial mechanical loads. Following data acquisition, movement trajectories were recorded at a high sampling rate of 200 Hz using a professional motion capture system. To ensure signal fidelity and remove high-frequency noise inherent in optical tracking, raw data were processed using a zero-lag Butterworth low-pass filter with a 6 Hz cutoff frequency. Finally, joint angles (*q* = [*θ*_*s*_, *θ*_*e*_]) were derived through inverse kinematics based on precise anthropometric measurements of each participant. This meticulous dataset provided the exact initial and final boundary conditions (*q*_*start*_, *q*_*end*_) required to benchmark the GJERK model’s predictions against the observed naturalistic human behavior.

### 4. Biomechanical Modeling of the Human Arm

For modeling the human arm, it was considered as a mathematical-mechanical robot model. Specifically, the arm was represented as a two-degree-of-freedom (2-DOF) planar manipulator operating in the parasagittal plane, consisting of two rigid links representing the upper arm and the forearm. The standard Lagrangian dynamics were employed to formulate the equations of motion as follows:

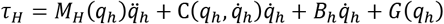

In this formulation, *q*_*h*_ = [*q*_*s*_, *q*_*e*_] denotes the vector of joint angles for the shoulder and elbow, whil 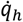 and 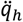 represent the corresponding joint velocities and accelerations, respectively. The matrices are defined as:

- *M*_*H*_(*q*_*h*_): is the symmetric, positive-definite inertia matrix.
- 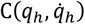: represents the Coriolis and centrifugal force vector.
- *G*(*q*_*h*_): is the gravitational torque vector, which accounts for the influence of the Earth’s gravity on each segment of the arm.
- *τ*_*H*_: represents the vector of joint torques exerted by the muscles.

This standard rigid-body dynamics model provides the necessary framework for integrating our gravity-aware cost function. Unlike models that treat dynamics as an external disturbance, our approach explicitly incorporates *G*(*q*_*h*_)within the optimization objective, thereby ensuring that the predicted movement trajectories are physically consistent with the gravitational field experienced by the participant throughout the reach.

### 5. Optimal Control Problem

#### 5.1 Formulation of the Optimal Control Problem for the Human Arm

To quantitatively investigate the principles of human motor planning, we model the Central Nervous System (CNS) as an optimal controller. The optimal trajectory planning problem for a general control system is first formulated as a Bolza-type optimal control problem, which provides a rigorous mathematical foundation for our analysis:

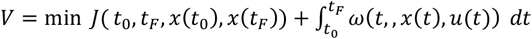

Where:

*t*_0_: the initial time, *t*_*f*_: at the final time.

*J*( *t*_0_, *t*_*f*_, *x*(*t*_0_), *x*(*t*_*f*_)) *dt*: The cost function at the boundary values, also called the Mayer term. In this case of the human arm model, it is equal to zero as the optimization focuses on path trajectory rather than terminal state costs.

*ω* (*t*,, *x*(*t*), *u*(*t*)) *dt*: The cost function along the trajectory, also called the Lagrange term. In this case of the human arm model, it is equal to one of the functions defined in Table 1.

Subject to the following constraints:

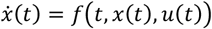: System dynamics; in this case of the human arm model, it is governed by 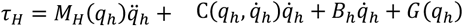, Where the state vector is 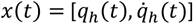 and the control input is *u*(*t*) = *τ*_*H*_(*t*).

*h*((*t, x*(*t*), *u*(*t*)) ≤ 0: Linear and nonlinear path constraints; in this case of the human arm model, in this case of the human arm model, these constraints link the state vector 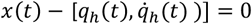, and the control input *u*(*t*) − *τ*_*H*_(*t*) =0, to the joint physical variables.

*g* (*t*_0_, *t*_*f*_, *x*(*t*_0_), *x*(*t*_*f*_)) ≤ 0: Bounds on state variables along the trajectory, these constraints link the end state vector 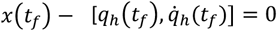, to the joint physical limits.

*x*_*low*_ ≤ *x*(*t*) ≤ *x*_*upp*_: Bounds on control inputs along the trajectory. in this case of the human arm model, these are defined by, −180 ≤ *q*_*s*_(*t*) ≤ 90, 0 ≤ *q*_*e*_(*t*) ≤ 160.

*x*_,*low*_ ≤ *x*(*t*_0_) ≤ *x*_,*upp*_: Boundary conditions on state variables at the initial time, these are defined by: 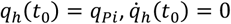

*x*_*low*_ ≤ *x*(*t*_*f*_) ≤ *x*_*f,upp*_ : Boundary conditions on state variables at the final time, these are defined by: *X*(*tf*)=0.85* *L*_*Total*_,*Y*(*tf*)=*free*.

### 5.2 Solving Optimal Control Problem for the Human Arm

To solve this problem, we transform it from its original continuous-time dynamic framework into a static algebraic representation, which allows the transformation of the system of differential equations into a Nonlinear Programming (NLP) problem solvable via numerical optimization algorithms like fmincon in MATLAB. We begin the discretization by partitioning the time interval [*t*_0_, *t*_*f*_] into N+1 nodes: *t* → *t*_0_ … *t*_*k*_ … *t*_*N*_, mapping the continuous state *x*(*t*), and control *u*(*t*) to discrete sequences *x*_0_ … *x*_*k*_ … *x*_*N*_ and *u*_0_ … *u*_*k*_ … *u*_*N*_. In this case of the human arm model, we utilized the Dynamics-Constrained Simpson method to discretize the optimal control equations above and transform them into a static algebraic form:[7]-[9], [32]-[39].

The mathematical equations derived using our method, without deductions, are given as follows:

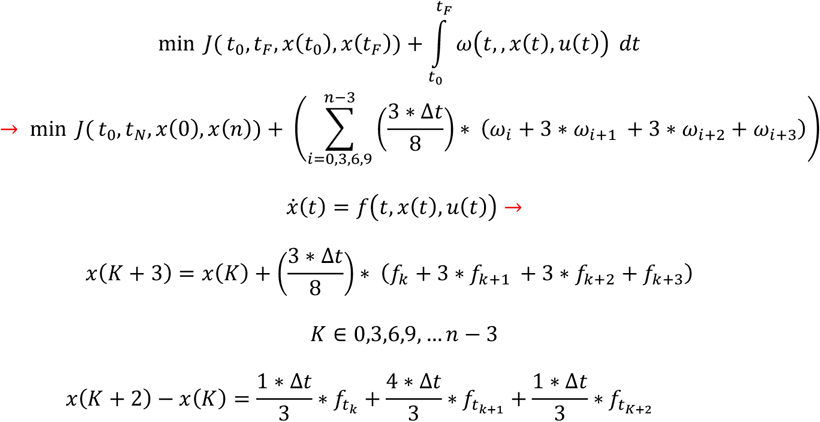

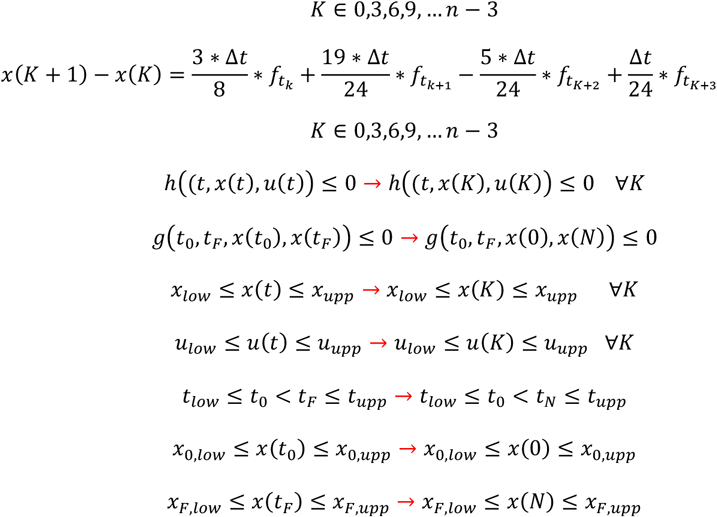

After discretization, the static optimization problem was solved using the fmincon function in MATLAB, we perform then cubic spline interpolation to reconstruct the continuous trajectories. This step is essential to recover the temporal evolution of the joint angles, velocities, and torques between the discrete nodes. By applying interpolation, we transform the solution from a set of discrete optimization variables into smooth, continuous functions.

### 6. Errors and affiliation transactions

### 6.1. linear speeds Errors

To evaluate the performance of the proposed cost functions, the linear velocity profiles were compared against experimental data. This process involved three main steps: normalization, area calculation via the trapezoidal rule, and membership mapping.

#### 6.1.1. Area Difference Calculation (Trapezoidal Rule)

The discrepancy between the simulated velocity (*V*_*model*_) and the experimental velocity (*V*_*data,j*_) was quantified by calculating the absolute area between the two curves. Since the velocity data is sampled at discrete time intervals(Δ*t*), the Trapezoidal Rule was employed for numerical integration:

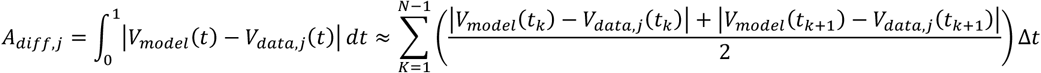

Where:

- N is the total number of samples in the normalized time interval [0,1].
- Δ*t* is the time step size.

#### 6.1.2. Membership Coefficient Mapping

The calculated area difference (*A*_*diff,j*_) was then transformed into a Membership Coefficient (*μ*_*j*_) using an exponential decay function. This coefficient represents the “degree of belonging” or similarity, where a value of 1 indicates an identical match:

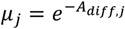

The final values reported in Table 2 are the arithmetic means of the membership coefficients calculated over 20 iterations for each initial position *P*_*i*_ :

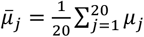

**Table 2:** The coefficients of belonging to the speeds of the cost functins in relation to the real speeds of the human arm.

| Cost Function | P1 | P2 | P3 | P4 | P5 | Average |
| --- | --- | --- | --- | --- | --- | --- |
| Gjerk (S1) | 0.8950 | 0.8603 | 0.9209 | 0.9099 | 0.8948 | 0.8933 |
| Gjerk (S2) | 0.8576 | 0.8838 | 0.8885 | 0.8813 | 0.8989 | 0.8820 |
| GJerk (S3) | 0.8652 | 0.9115 | 0.8758 | 0.8334 | 0.8758 | 0.8723 |
| Gjerk (S4) | 0.8600 | 0.9044 | 0.9056 | 0.9019 | 0.8720 | 0.8887 |
| GJerk (S5) | 0.8959 | 0.8625 | 0.9090 | 0.8220 | 0.9079 | 0.8794 |
| GJerk (S6) | 0.8977 | 0.9028 | 0.8932 | 0.8950 | 0.8930 | 0.8963 |
| **AverageG** | **0.8785** | **0.8852** | **0.8988** | **0.8739** | **0.8904** | **0.88533** |
| Energy +AJerk (S1) | 0.9035 | 0.8796 | 0.9061 | 0.8545 | 0.8993 | 0.8886 |
| Energy +AJerk (S2) | 0.8949 | 0.8726 | 0.8872 | 0.8480 | 0.8960 | 0.8797 |
| Energy +AJerk (S3) | 0.8488 | 0.8700 | 0.8585 | 0.7983 | 0.8477 | 0.8446 |
| Energy +AJerk (S4) | 0.8435 | 0.8921 | 0.9127 | 0.8705 | 0.8249 | 0.8687 |
| Energy +AJerk (S5) | 0.8935 | 0.8412 | 0.9120 | 0.7880 | 0.9040 | 0.8677 |
| Energy +AJerk (S6) | 0.8860 | 0.9002 | 0.8980 | 0.8827 | 0.9119 | 0.8957 |
| **AverageE+AJ** | **0.8784** | **0.8760** | **0.8957** | **0.8403** | **0.8806** | **0.87417** |
| Energy+AACC (S1) | 0.8074 | 0.8408 | 0.8630 | 0.8209 | 0.7944 | 0.8253 |
| Energy +AACC (S2) | 0.7892 | 0.8197 | 0.8485 | 0.8133 | 0.8033 | 0.8148 |
| Energy +AACC (S3) | 0.7552 | 0.8317 | 0.7515 | 0.7810 | 0.7674 | 0.7736 |
| Energy +AACC (S4) | 0.7549 | 0.8164 | 0.8333 | 0.8104 | 0.7269 | 0.7883 |
| Energy +AACC (S5) | 0.7899 | 0.8200 | 0.8704 | 0.7350 | 0.7960 | 0.8022 |
| Energy +AACC (S6) | 0.7807 | 0.8494 | 0.8734 | 0.8139 | 0.7883 | 0.8211 |
| **AverageE+AC** | **0.7795** | **0.8296** | **0.8400** | **0.7957** | **0.7793** | **0.80422** |

Furthermore, Table 3 also presents the standard deviation of these average membership coefficients, providing insight into the variability and consistency of each cost function in reproducing human-arm speeds.

**Table 3:** Standard Deviation of Average Speed-Matching Coefficients for All Cost Functions.

| Cost Function | Standard Deviation (STD) |
| --- | --- |
| Gjerk | 0.0083 |
| Energy + AJerk | 0.0166 |
| Energy + AACC | 0.0184 |

### 6.2. Path Accuracy and Error Quantification

To evaluate the performance of the proposed cost functions, the deviation between the simulated trajectories and the experimental (measured) data was quantified using two distinct metrics: Cartesian error and curvature error as illustrated in Figure 3. These metrics ensure a comprehensive assessment of both spatial positioning and path geometry.

**Figure 3:**
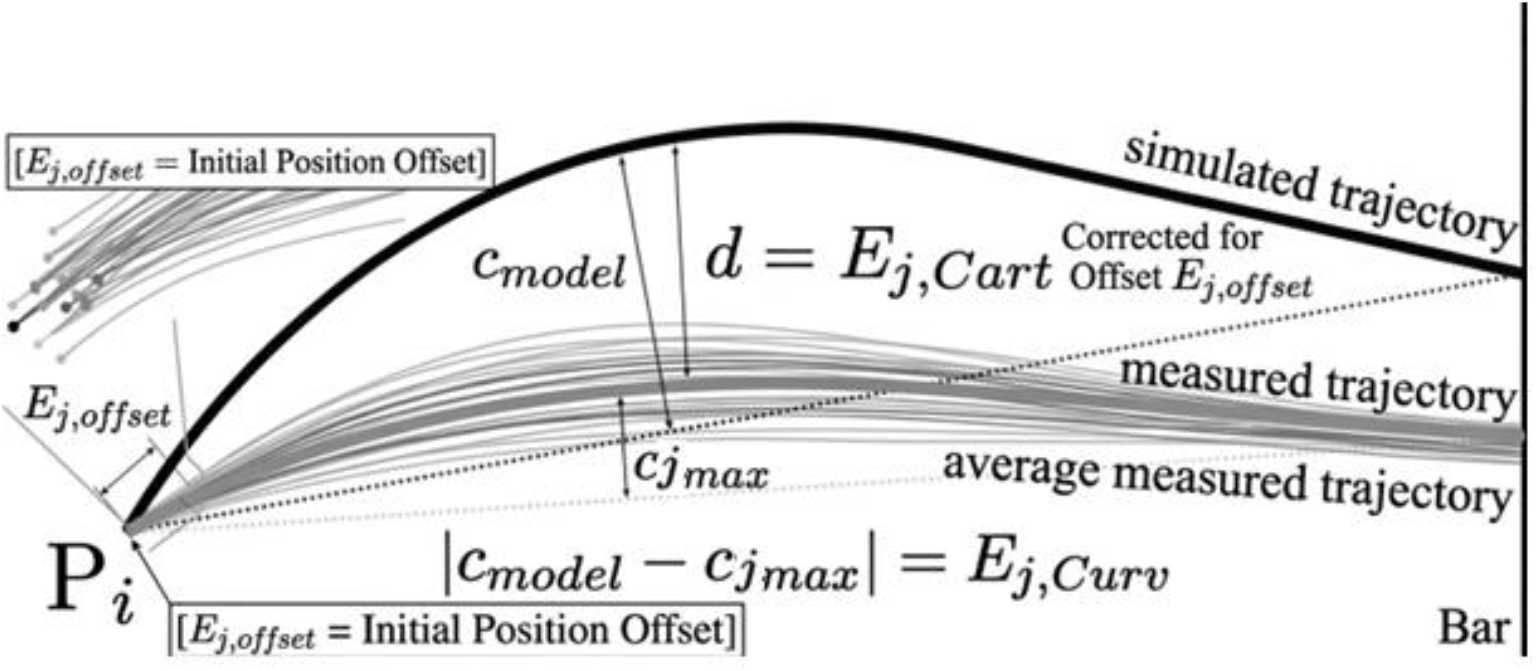
Statistical methods for calculating mathematical errors of paths

**Figure 3:**
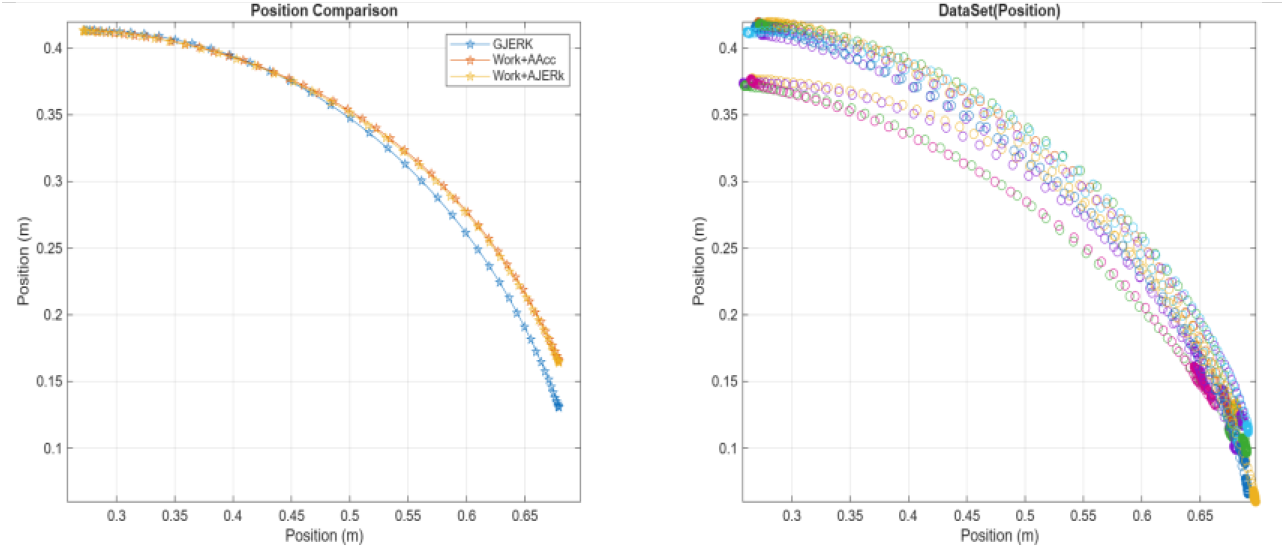
The following figure illustrates the mathematical function trajectories on the left and the actual experimental data trajectories of real human arm movement on the right, during the execution of the first transition P1 by subject S1

#### 6.2.1. Cartesian Error (*E*_*i*,*Card*_)

The Cartesian error represents the direct spatial deviation between the predicted and actual paths. To account for potential human error during the initial positioning of the limb, an offset correction was applied:

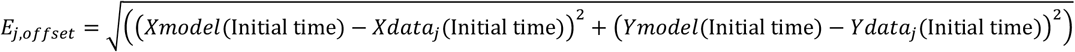

Where (*Xmodel, Ymodel*) and (*Xdata, Ydata*) are the coordinates of the model. The mean Cartesian error for each point across N=20 repetitions is then calculated as:

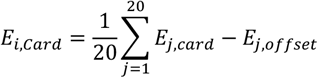

#### 6.2.2 Curvature Error (*E*_*i*,*curv*_)

To capture the differences in the “shape” or “convexity” of the motion, the curvature error was defined. While the computational model produces a single deterministic path (*C*_*model*_), the human data varies across trials (*C*_*data,j*_). The curvature error is the mean absolute difference between these values:

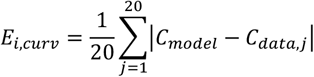

#### 6.2.3. Total Error (*E*_*i*,*total*_)

The overall performance for each initial position *P*_*i*_ is expressed as the sum of these two components. This combined metric ensures that a “good” model must match both the position and the biological “style” of the movement:

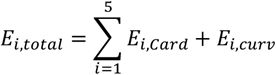

Table 3 presents the total error values for each point, providing a clear summary of both spatial deviation and curvature variation.

Furthermore, Table 5 also presents the standard deviation of these average membership coefficients, providing insight into the variability and consistency of each cost function in reproducing human-arm positions.

## 7. Practical results

### 7.1. linear speeds

The empirical superiority of the proposed GJERK model, characterized by an overall mean Membership Coefficient of 0.88533, provides strong validation for the shift from static penalty-based frameworks to gravity-weighted dynamic operators. Theoretically, the performance stratification observed in Table 2—where GJERK consistently outperforms the baseline Energy+AJERK (0.87417) and Energy+AACC (0.80422) models—is a direct consequence of how each framework reconciles the mechanical implications of the gravitational torque sign(G_i_(θ)) Specifically, the model’s foundation on continuous softplus-based modulation, 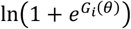 allows it to dynamically reallocate ‘jerk variance’ to preferred actuators; this enables the GJERK architecture to maintain high consistency across all test positions (P1–P5), whereas static models experience accuracy degradation due to the destabilizing effect of negative weights in assistive gravitational configurations. Furthermore, the superior vertical mean of the GJERK model provides quantitative evidence for emergent biological behaviors, such as joint ‘freezing’ and the stabilization of the vertebral column, confirming that these patterns arise naturally from jerk minimization within a high-potential-energy landscape rather than active neural restriction. By successfully integrating the physical modulation of gravitational torque into the cost function, the GJERK model effectively demonstrates that rhythmic motor patterns, such as arm swinging, emerge as a consequence of optimizing jerk distribution to balance postural integrity and energetic costs across all vertical movement trajectories, thereby validating the efficacy of the softplus operator in capturing the full complexity of human musculoskeletal dynamics. In addition, the standard deviation analysis presented in Table 3 further reinforces the stability of the GJERK model; its markedly lower STD (0.0083) compared to Energy+AJerk (0.0166) and Energy+AACC (0.0184) demonstrates that the GJERK framework produces highly consistent speed-matching coefficients with minimal variability, a property essential for accurately characterizing physical laws and ensuring reproducible biomechanical behavior.

### 7.2. Path Accuracy

The results presented in Table 4 demonstrate that the GJERK model provides a more precise geometric characterization of human arm movement, achieving a minimum total mean error of 12.49 cm, compared to 14.02 cm for the Work+AJERK model and 13.91 cm for the Work+AACC model. This lower error margin quantifies the model’s superior ability to capture the specific “shape” (curvature) and spatial positioning of biological trajectories. While models relying on static weighting often produce unnatural deviations and sharp geometric distortions inconsistent with human motor style, the GJERK model maintains path fluidity across all test positions (P1–P5).

**Table 4:**
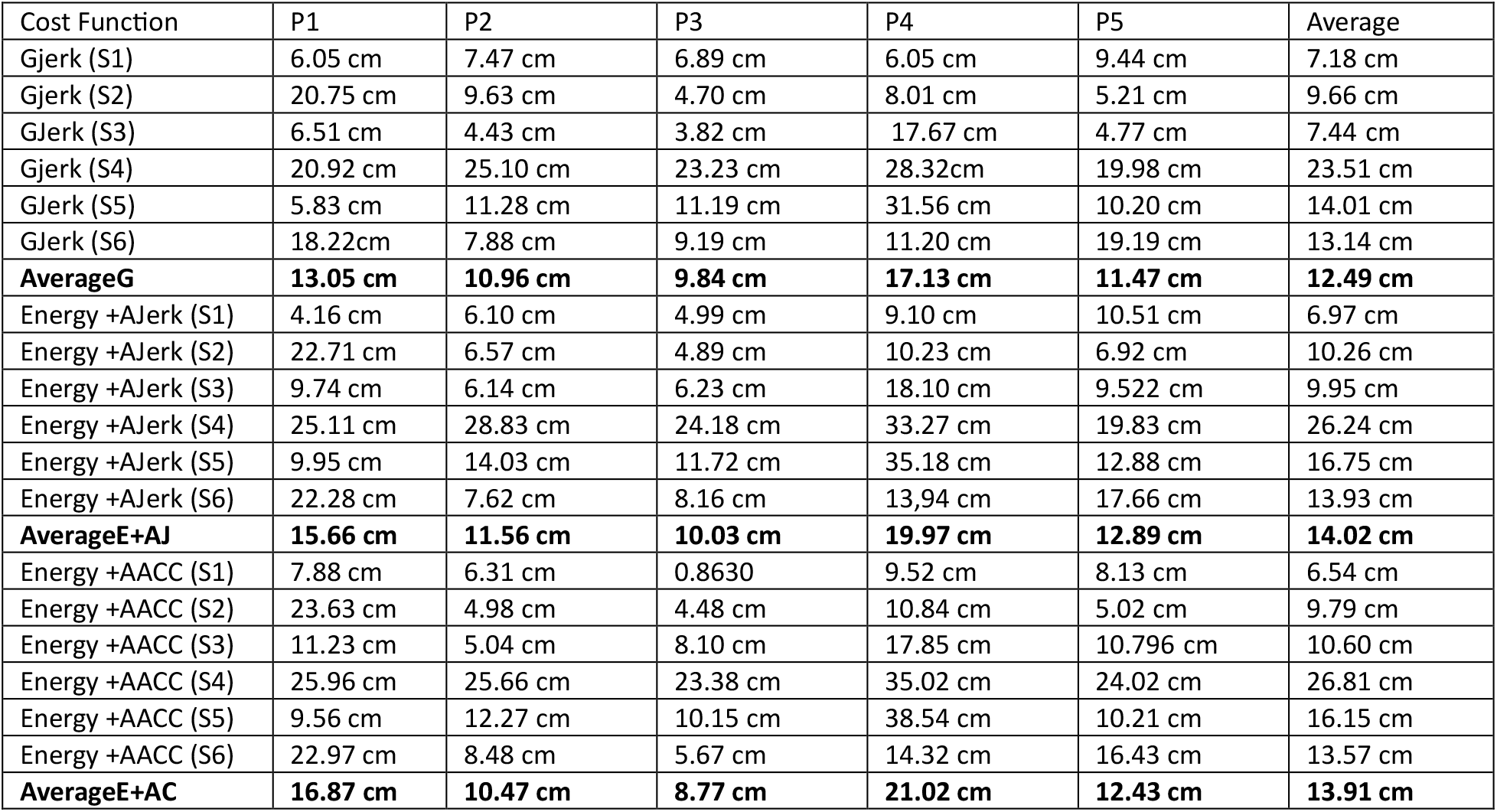
Error values (Etotal) in terms of cost functions and initial positions.

Additionally, the low standard deviation reported in Table 5 further confirms that our GJERK-based cost function provides superior stability, as its trajectories exhibit minimal positional variance compared to all other models, which is essential for describing physical laws.

**Table 5:** Standard Deviation of Average positions-Matching Coefficients for All Cost Functions.

| Cost Function | Standard Deviation (STD) |
| --- | --- |
| Gjerk | 1.399 cm |
| Energy + AJerk | 2.724 cm |
| Energy + AACC | 3.371 cm |

This accurate geometric characterization reveals how the model avoids the kinematic artifacts imposed by static constraints; by employing a dynamic mechanism that aligns the proposed path with gravitational physics rather than opposing it, the resulting trajectory faithfully describes the innate “motor style” of the human arm with high fidelity and minimal deviation from experimental reality.

### 7.3. Visual Validation

In addition to the quantitative analysis presented earlier, the strength of our study is further reinforced by incorporating visual evidence that enhances confidence in the validity of the obtained motion dynamics. While the numerical indicators clearly demonstrate the superiority of the **GJERK** model, these values alone cannot fully convey the qualitative nature of the resulting movement trajectories. Therefore, this section establishes a direct and coherent link between the statistical outcomes and the detailed visual representations corresponding to each of the five movement cases, which are comprehensively illustrated in **Figures 3–32**. These visual demonstrations provide concrete proof of the distinctive characteristics related to both the overall “shape” of the trajectory and the inter-joint “coordination” patterns, thereby offering a clearer understanding of how our proposed model excels in reproducing natural human motion. Figures 3–32 present the experimental results for a randomly selected participant performing the five movements, serving as representative examples that visually confirm the dynamic behavior captured by our approach.

In the first case, we observe from Figures 3, 5, and 7 that the cost function proposed by us (GJERK)—shown on the left side of each figure—resembles the actual trajectories of human arm motion more closely during the execution of movement P1 for subjects 1, 2, and 3. In contrast to the other cost functions (Energy+Acc and Energy+Ajerk), our proposed function exhibits a more pronounced inward curvature, and its final stopping point lies closer to the end points of the real human arm trajectories. Furthermore, the velocity profile associated with our proposed cost function GJERK demonstrates an asymmetric bell shaped behavior, reaching its maximum at a time of 0.43, as illustrated in Figures 4, 6, and 8 during movement P1. This value is approximately equal to the average peak reaching time observed across multiple subjects performing the same movement. Meanwhile, the remaining cost functions Energy +Acc and Energy +Ajerk fail to reproduce this characteristic behavior, as they neither generate an asymmetric bell shaped velocity profile nor approximate the experimentally observed average peak reaching time for movement P1.

**Figure 4:**
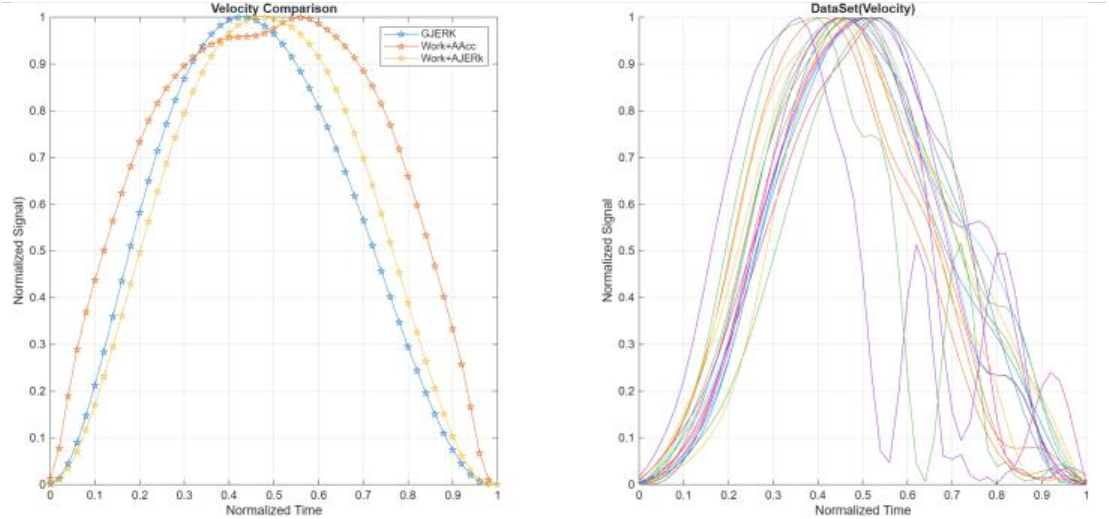
The following figure illustrates the velocity profiles for the mathematical functions on the left and the actual velocities of human arm movement on the right, during the execution of transition P1 by subject S1.

**Figure 5:**
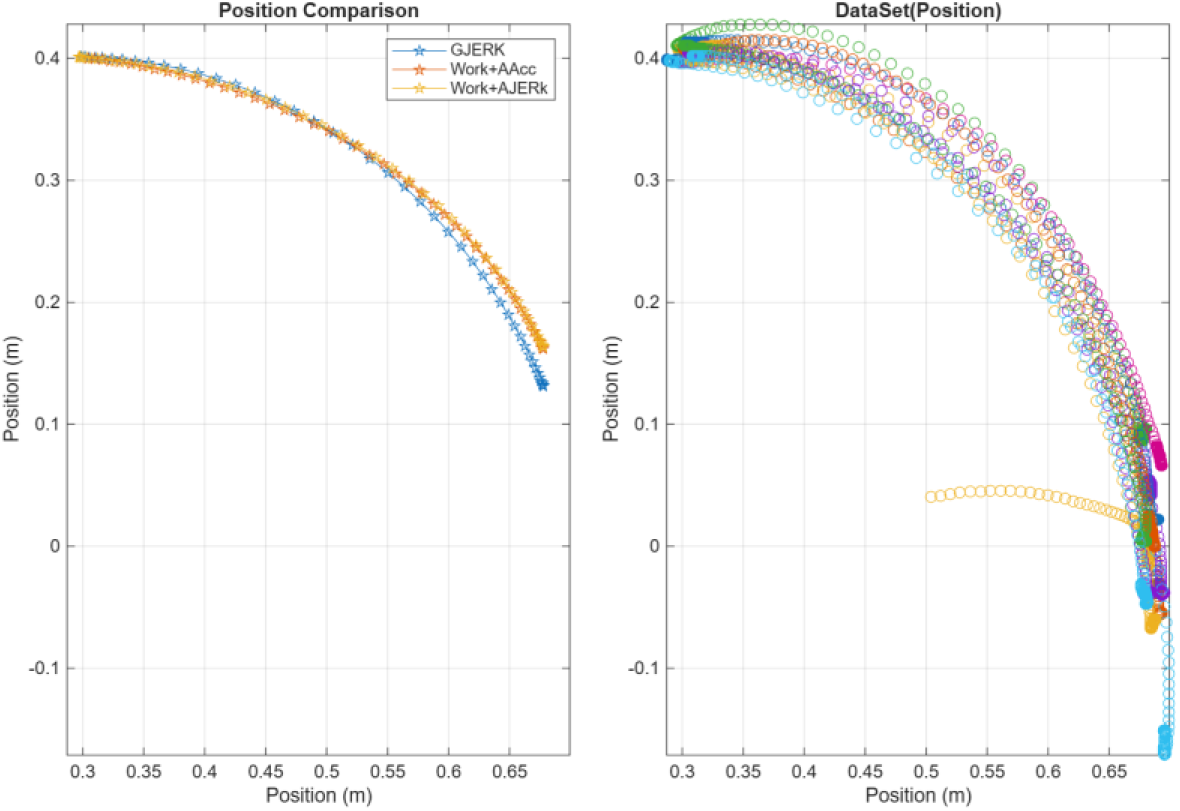
The following figure illustrates the mathematical function trajectories on the left and the actual experimental data trajectories of real human arm movement on the right, during the execution of the first transition P1 by subject S2

**Figure 6:**
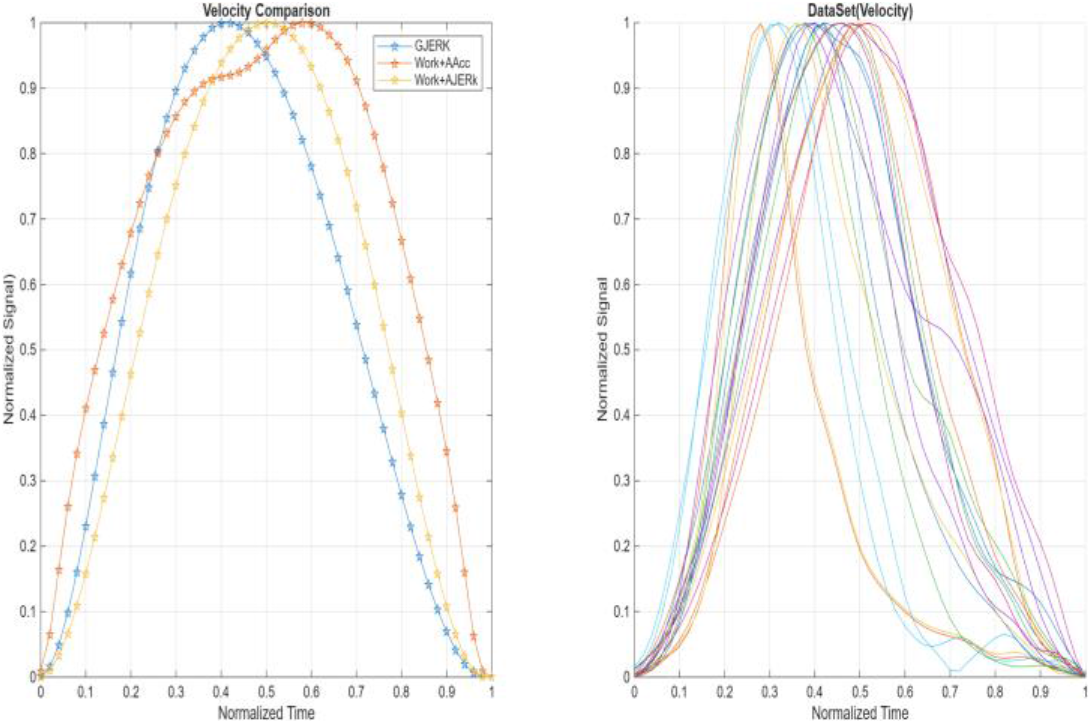
The following figure illustrates the velocity profiles for the mathematical functions on the left and the actual velocities of human arm movement on the right, during the execution of transition P1 by subject S2.

**Figure 7:**
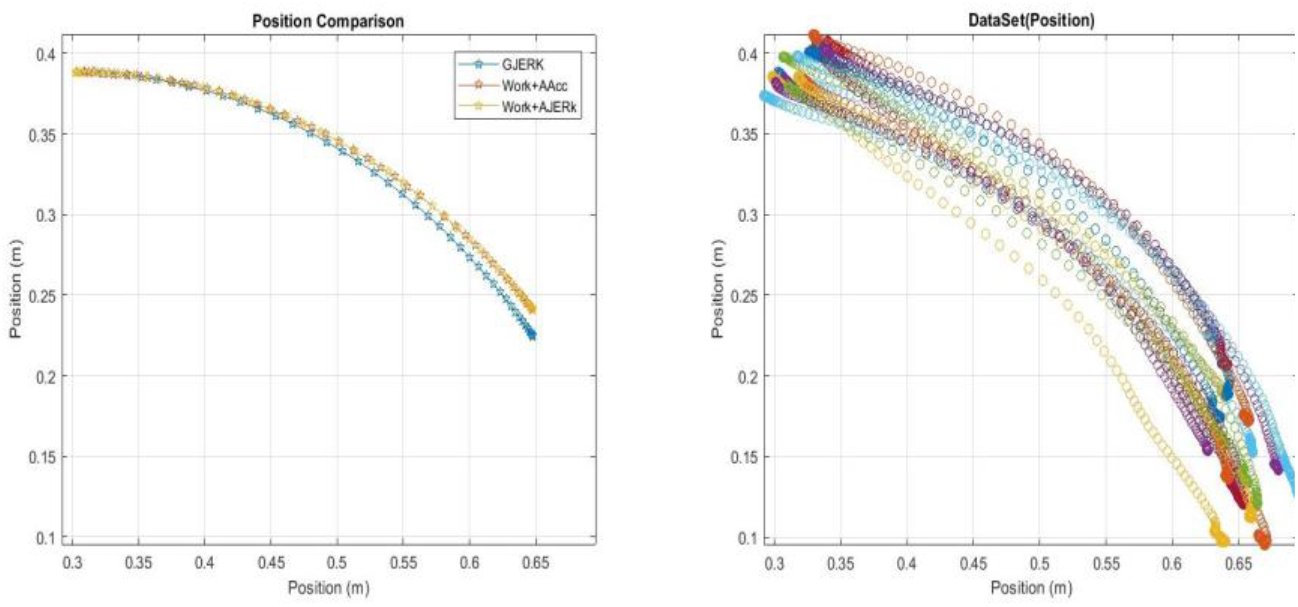
The following figure illustrates the mathematical function trajectories on the left and the actual experimental data trajectories of real human arm movement on the right, during the execution of the first transition P1 by subject S3

**Figure 8:**
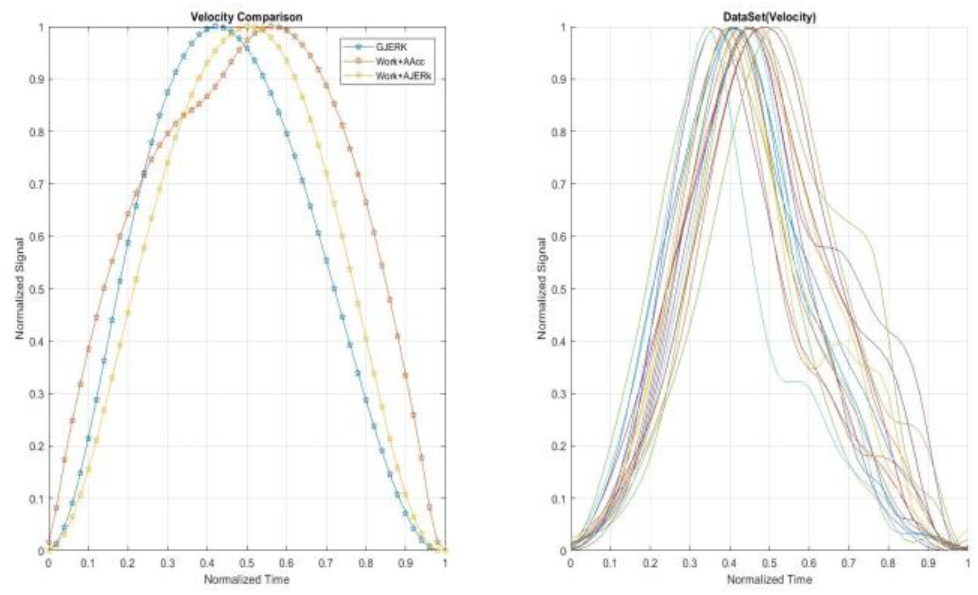
The following figure illustrates the velocity profiles for the mathematical functions on the left and the actual velocities of human arm movement on the right, during the execution of transition P1 by subject S3.

In the second case, We observe from Figures 9, 11, and 13 that our proposed cost function (GJERK)—displayed on the left side of each figure—captures the essential characteristics of the human arm trajectories during the execution of movement P2 for subjects 1, 2, and 3. In contrast to movement P1, this transition constitutes a highly sensitive and variable case, as the spatial trajectories differ markedly across individuals. Some subjects tend to move their fingertip toward a noticeably higher final position relative to the starting point, whereas others prefer end points that remain nearly level with the initial position or only slightly elevated. Despite this substantial inter subject variability, the trajectory generated by our proposed cost function GJERK lies close to the overall central trend of all participants. Although it does not replicate the exact path of any single subject, it provides a representative compromise that reflects the collective behavior of the group, thereby achieving a high degree of accuracy relative to the distribution of human trajectories. Furthermore, the velocity profiles associated with movement P2—as illustrated in Figures 10, 12, and 14—exhibit a distinctly irregular and strongly asymmetric structure across subjects. Humans naturally produce non symmetric velocity curves in this transition, indicating the absence of a consistent motor strategy for this movement. Notably, the experimental velocity curves consistently show a bell shaped peak occurring around 0.43, a characteristic that is also reproduced by the velocity profile generated by our proposed cost function GJERK. This agreement demonstrates that, even under conditions of pronounced spatial variability, our cost function successfully captures the fundamental temporal dynamics of human motion. In contrast, the remaining cost functions Energy +Acc and Energy +Ajerk fail to reproduce these features, as they neither generate realistic asymmetric velocity profiles nor adapt to the considerable variability inherent in movement P2. In the third case, we observe from Figures 15, 17, and 19 that the cost function proposed by us (GJERK)—shown on the left side of each figure—captures the overall structure of the human arm trajectories during the execution of movement P3 for subjects 1, 2, and 3. Unlike movement P1, this transition represents a particularly critical case, as the spatial trajectories vary substantially from one subject to another. While some individuals tend to reach a final fingertip position that is significantly higher than the initial point, others prefer end points that remain nearly at the same vertical level or only slightly elevated. Despite this pronounced inter subject variability, our proposed cost function GJERK generates a trajectory that lies near the central tendency of all subjects. Although the resulting path does not perfectly match any single participant, it provides a balanced compromise among the different observed behaviors, thereby achieving a high level of accuracy relative to the overall distribution of human trajectories. Moreover, the velocity profiles recorded for movement P3—as shown in Figures 16, 18, and 20—reveal a strongly irregular and noticeably asymmetric pattern across subjects. Humans naturally produce markedly non symmetric velocity curves in this transition, reflecting the absence of a unified motor strategy for this movement. Importantly, these experimental velocity curves consistently exhibit a bell shaped peak occurring around 0.43, a feature that is also reproduced by the velocity profile generated by our proposed cost function GJERK. Meanwhile, the remaining cost functions Work+Acc and Work+Ajerk fail to reproduce this behavior, as they neither generate realistic asymmetric velocity profiles nor adapt to the substantial variability inherent in movement P3. In the fourth case, we observe from Figures 21, 23, and 25 that the cost function proposed by us (GJERK)—shown on the left side of each figure—resembles the actual trajectories of human arm motion more closely during the execution of movement P4 for subjects 1, 2, and 3. In contrast to the other cost functions Work+Acc and Work+Ajerk, our proposed function exhibits a more pronounced inward curvature, and its final stopping point lies closer to the end points of the real human arm trajectories. Furthermore, the velocity profile associated with our proposed cost function GJERK demonstrates an asymmetric bell shaped behavior, reaching its maximum at a time of 0.43, as illustrated in Figures 22, 24, and 26 during movement P4. However, the experimental data show that the average peak reaching time across multiple subjects performing the same movement is approximately 0.40. Meanwhile, the remaining cost functions Work+Acc and Work+Ajerk fail to reproduce this characteristic behavior, as they neither generate an asymmetric bell shaped velocity profile nor approximate the experimentally observed average peak reaching time for movement P4. In the fifth and final case, We observe from Figures 27, 29, and 31 that our proposed cost function (GJERK)—displayed on the left side of each figure—offers a trajectory that more faithfully reflects the natural motion of the human arm during the execution of movement P5 for subjects 1, 2, and 3. Consistent with the observations made for movement P1, the trajectory generated by our cost function GJERK shows a clearer inward curvature, and its final stopping point is positioned closer to the experimentally measured end points of the actual human arm trajectories. In contrast, the alternative cost functions Energy +Acc and Energy +Ajerk introduce noticeable fluctuations and irregular deviations along the path, which do not correspond to realistic human movement patterns. Additionally, the velocity profile produced by our proposed cost function GJERK exhibits an asymmetric bell shaped form, reaching its peak at 0.41, as shown in Figures 28, 30, and 32 for movement P5. This peak reaching time closely matches the average value observed across multiple subjects performing the same movement. Meanwhile, the other cost functions Energy +Acc and Energy +Ajerk fail to replicate this characteristic behavior, as they neither generate an asymmetric bell shaped velocity curve nor approximate the experimentally observed average peak reaching time for movement P5.

**Figure 9:**
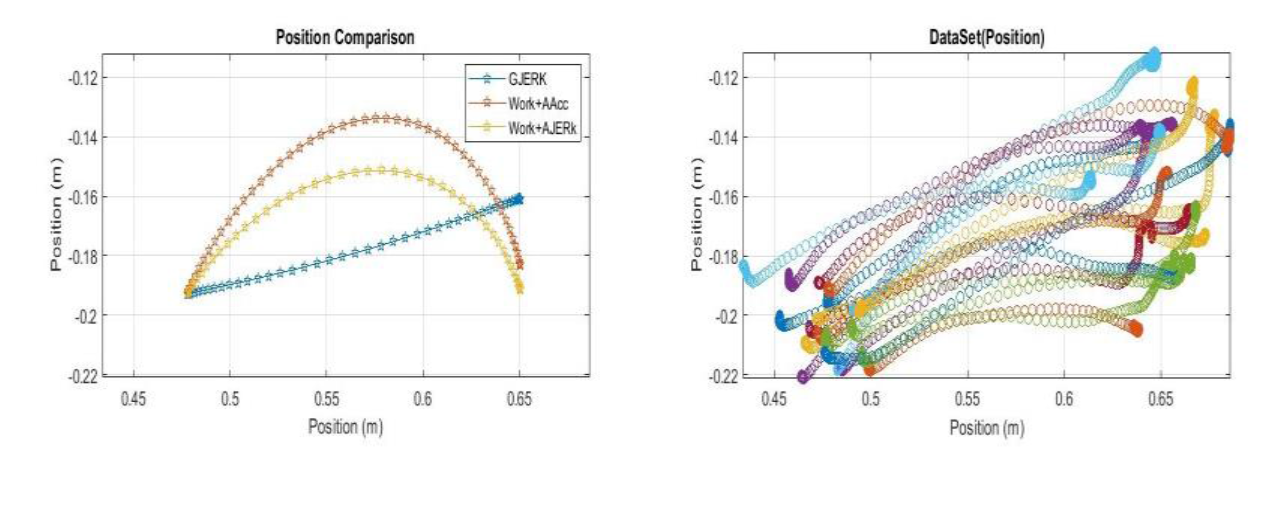
The following figure illustrates the Cost functions trajectories on the left and the actual experimental data trajectories of real human arm movement on the right, during the execution of the first transition P2 by subject S1

**Figure 10:**
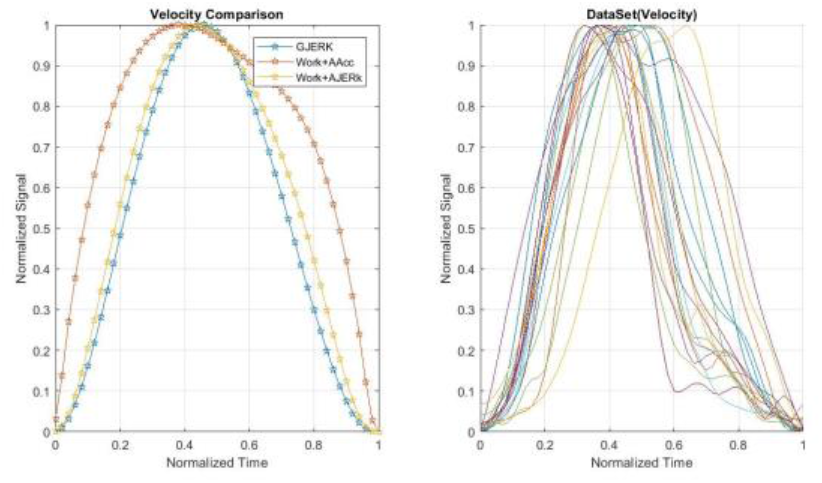
The following figure illustrates the velocity profiles for the Cost functions on the left and the actual velocities of human arm movement on the right, during the execution of transition P2 by subject S1.

**Figure 11:**
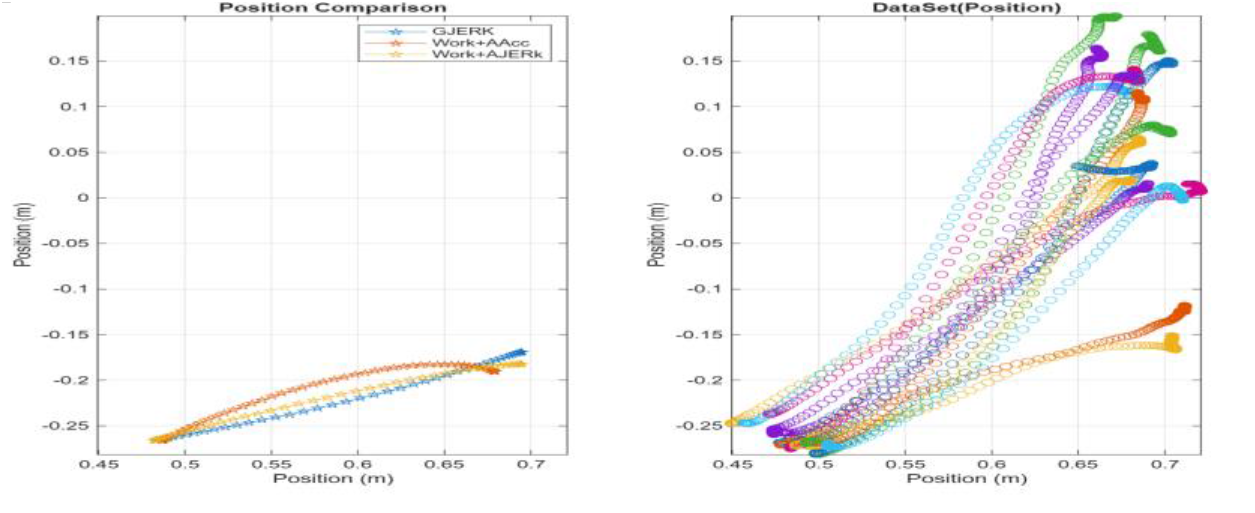
The following figure illustrates the Cost functions trajectories on the left and the actual experimental data trajectories of real human arm movement on the right, during the execution of the first transition P2 by subject S2.

**Figure 12:**
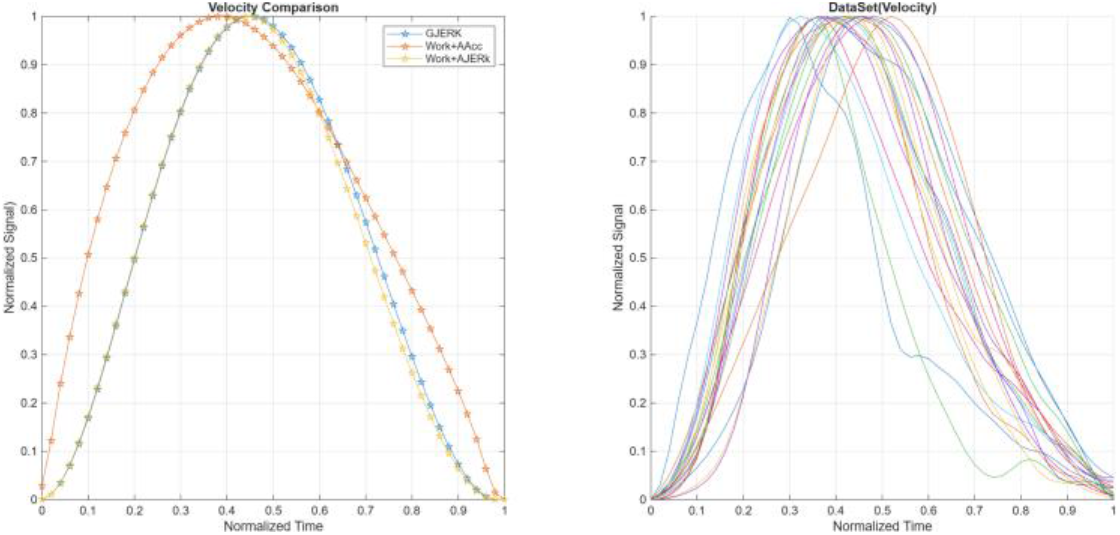
The following figure illustrates the velocity profiles for the Cost functions on the left and the actual velocities of human arm movement on the right, during the execution of transition P2 by subject S2.

**Figure 13:**
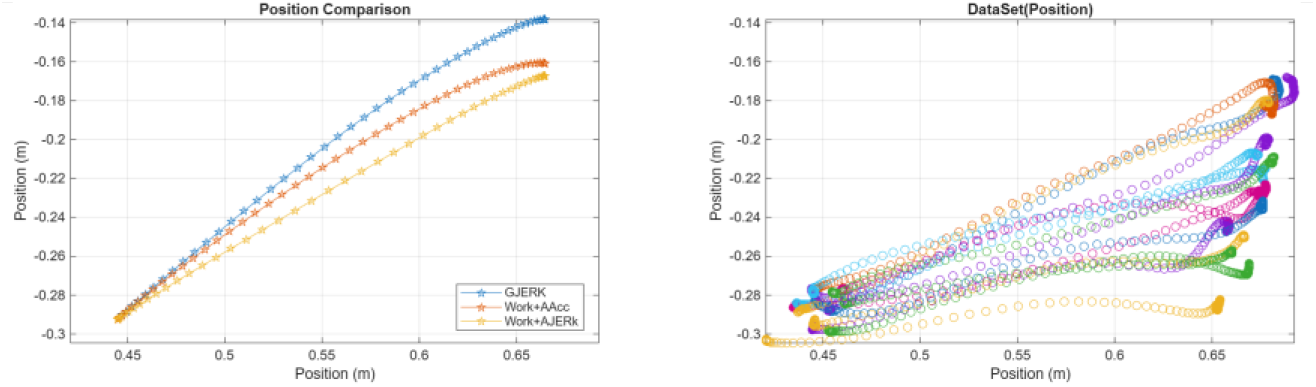
The following figure illustrates the Cost functions trajectories on the left and the actual experimental data trajectories of real human arm movement on the right, during the execution of the first transition P2 by subject S3.

**Figure 14:**
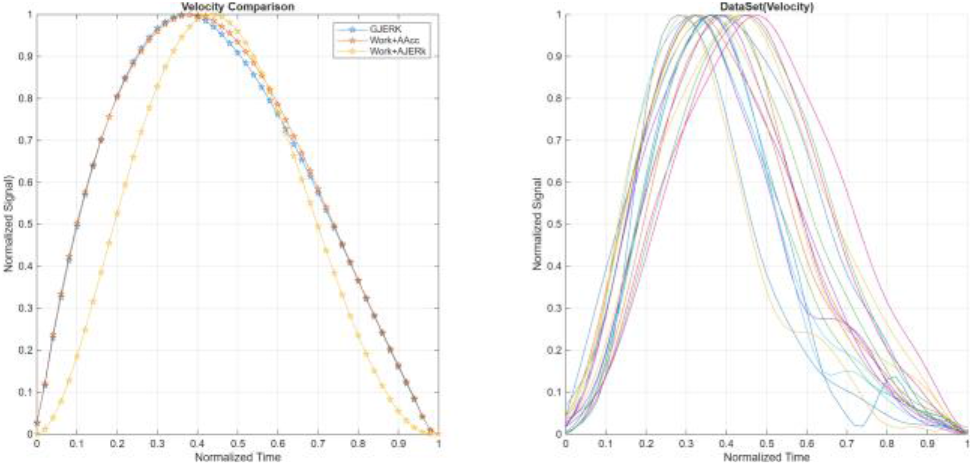
The following figure illustrates the velocity profiles for the Cost functions on the left and the actual velocities of human arm movement on the right, during the execution of transition P2 by subject S3

**Figure 15:**
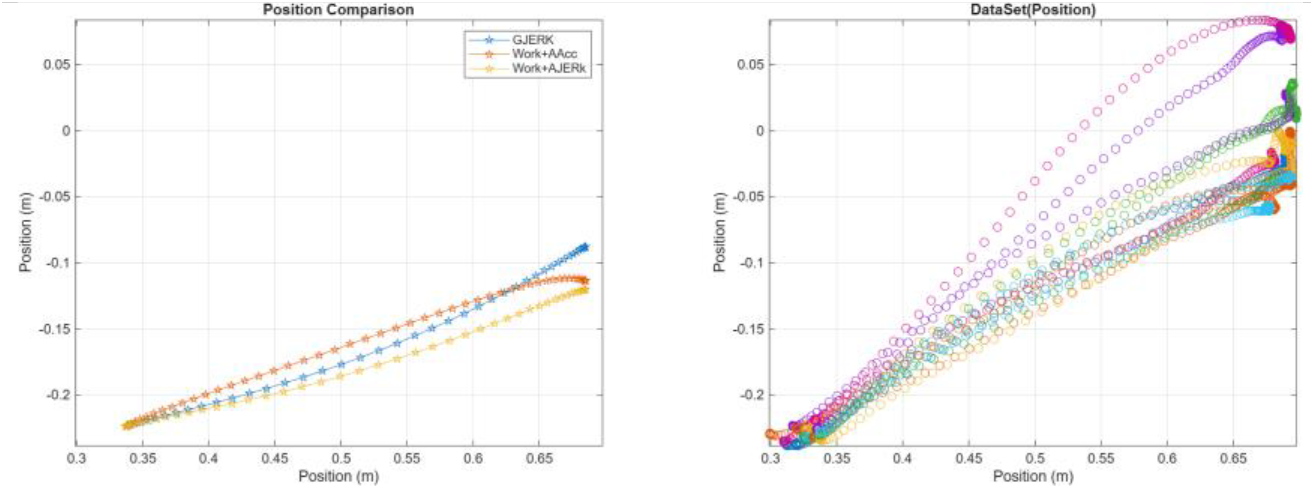
The following figure illustrates the Cost functions trajectories on the left and the actual experimental data trajectories of real human arm movement on the right, during the execution of the first transition P3 by subject S1

**Figure 16:**
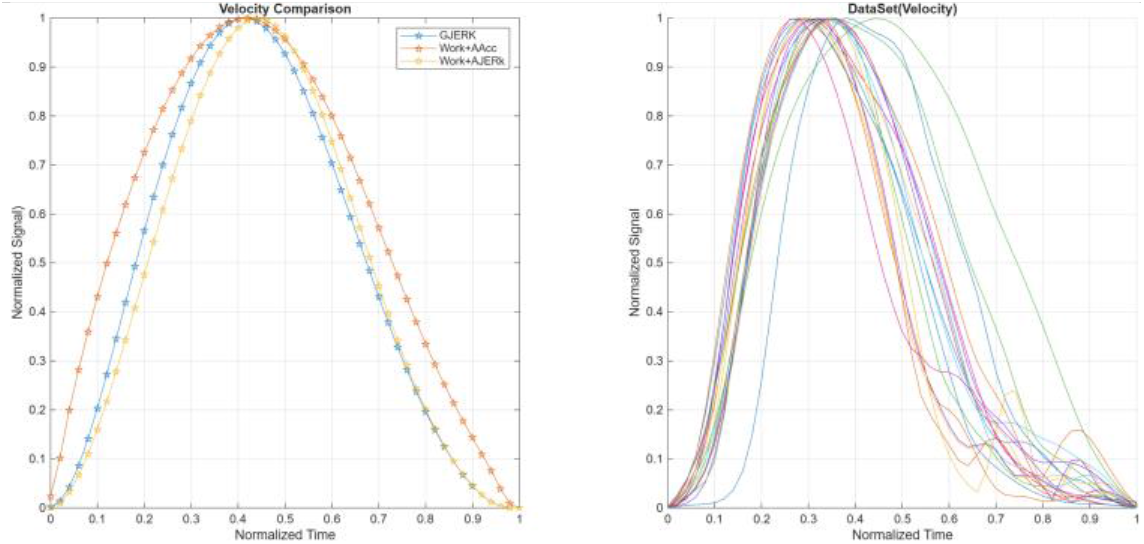
The following figure illustrates the velocity profiles for the Cost functions on the left and the actual velocities of human arm movement on the right, during the execution of transition P3 by subject S1.

**Figure 17:**
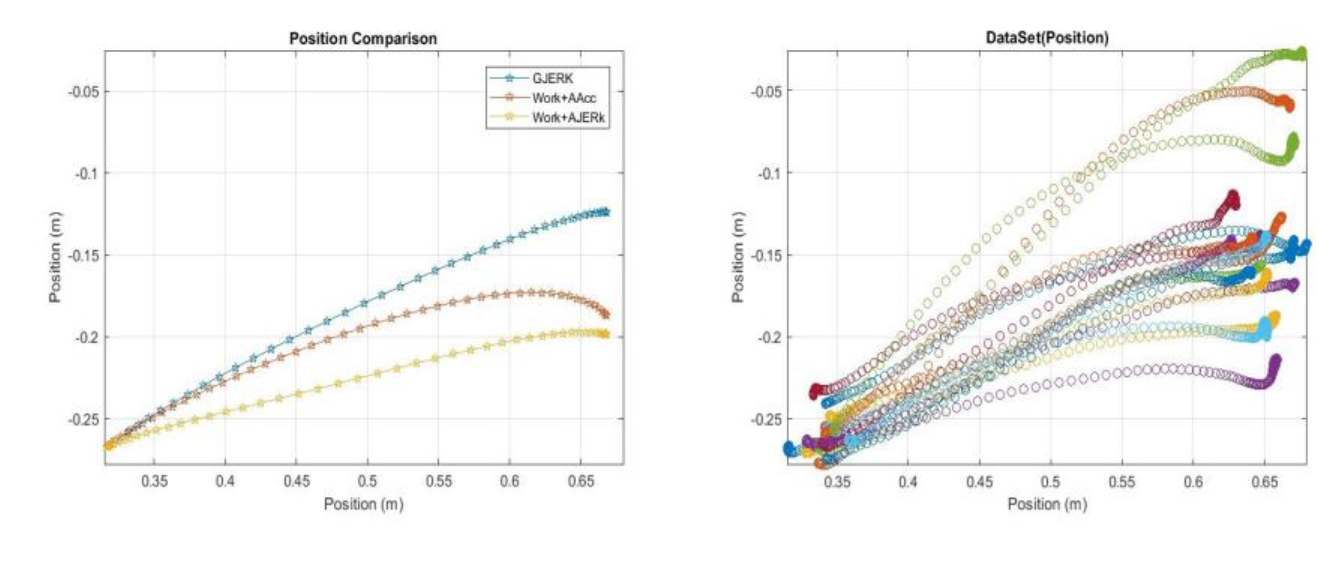
The following figure illustrates the Cost functions trajectories on the left and the actual experimental data trajectories of real human arm movement on the right, during the execution of the first transition P3 by subject S2

**Figure 18:**
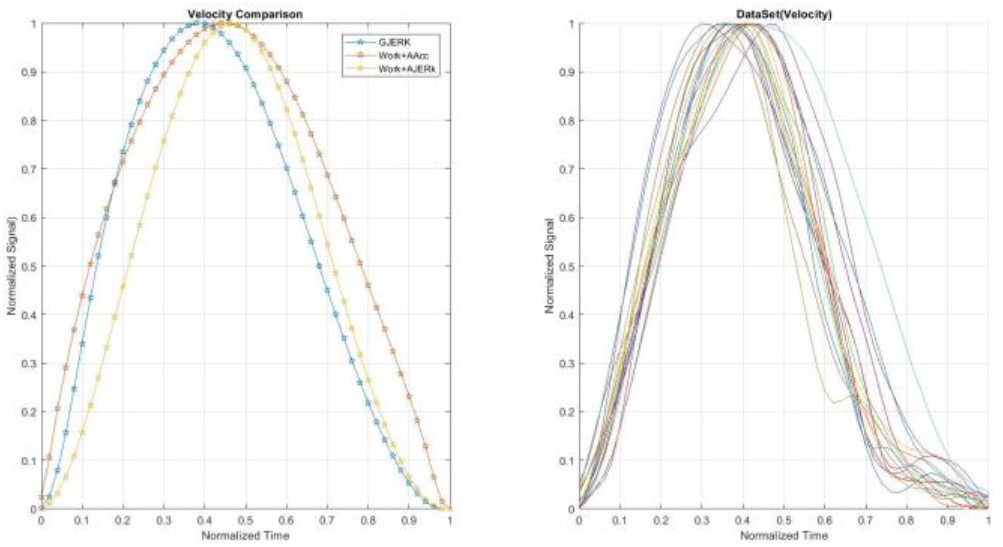
The following figure illustrates the velocity profiles for the Cost functions on the left and the actual velocities of human arm movement on the right, during the execution of transition P3 by subject S2.

**Figure 19:**
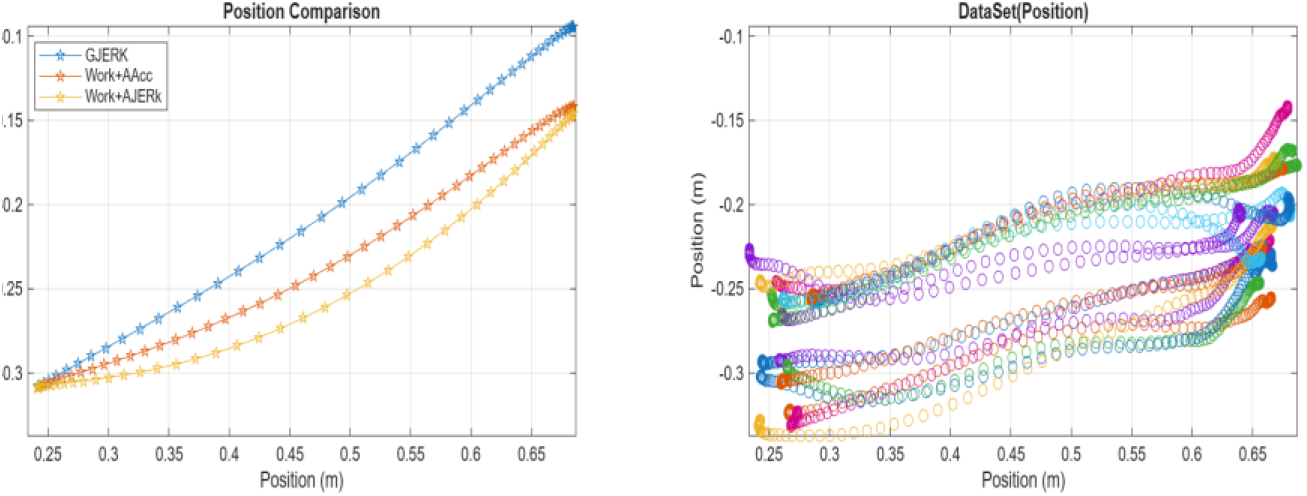
The following figure illustrates the Cost functions trajectories on the left and the actual experimental data trajectories of real human arm movement on the right, during the execution of the first transition P3 by subject S3

**Figure 20:**
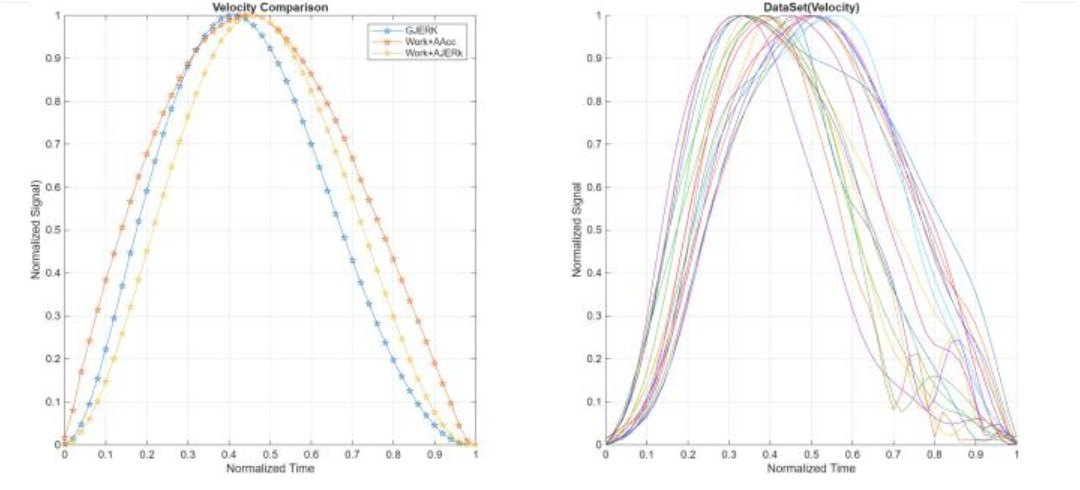
The following figure illustrates the velocity profiles for the Cost functions on the left and the actual velocities of human arm movement on the right, during the execution of transition P3 by subject S3.

**Figure 21:**
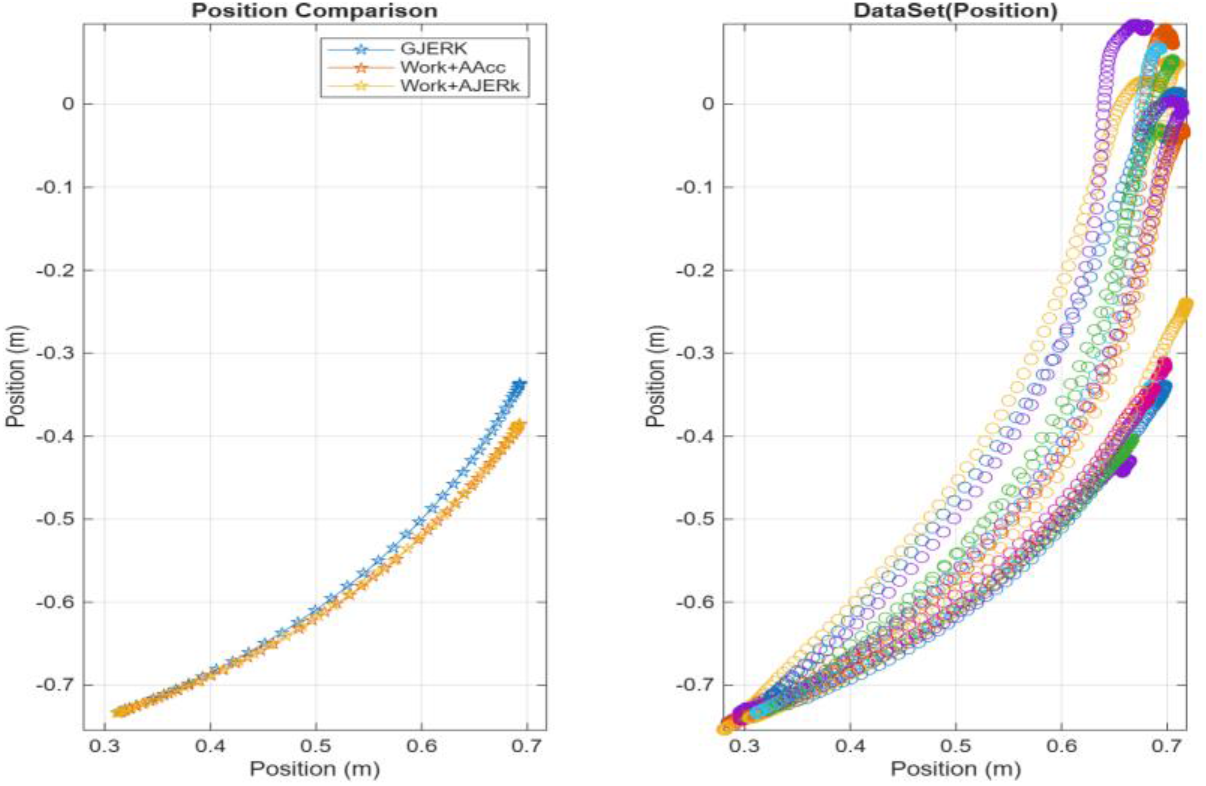
The following figure illustrates the Cost functions trajectories on the left and the actual experimental data trajectories of real human arm movement on the right, during the execution of the first transition P4 by subject S1.

**Figure 22:**
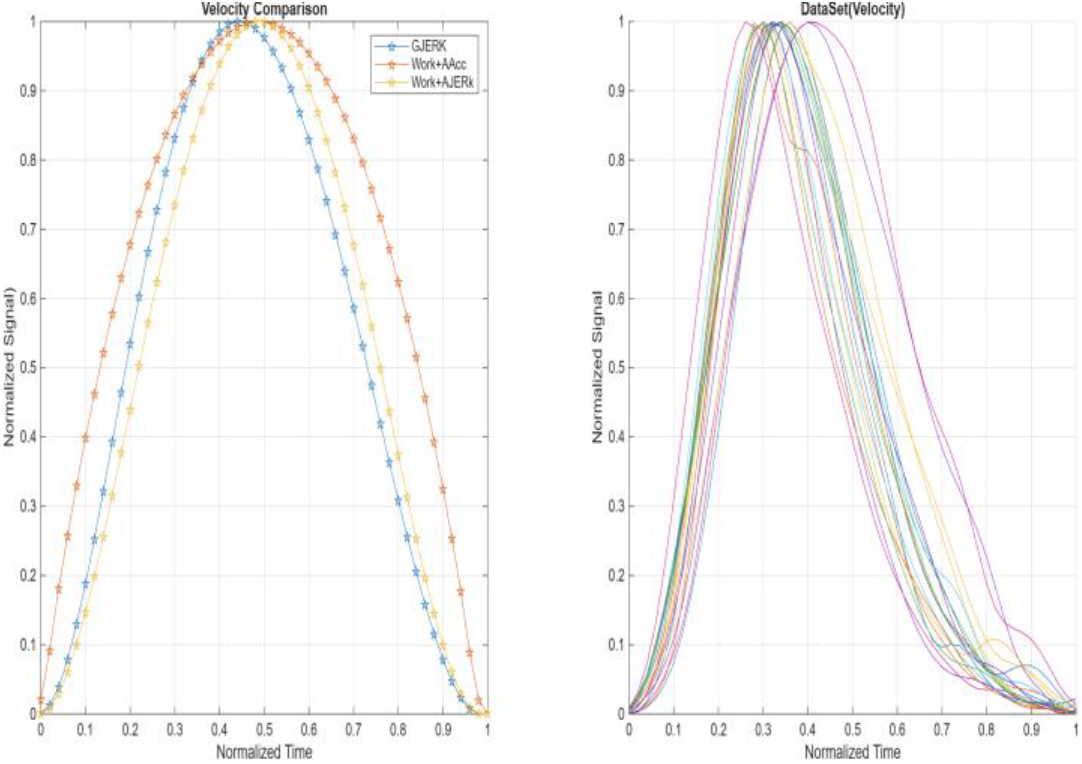
The following figure illustrates the velocity profiles for the Cost functions on the left and the actual velocities of human arm movement on the right, during the execution of transition P4 by subject S1.

**Figure 23:**
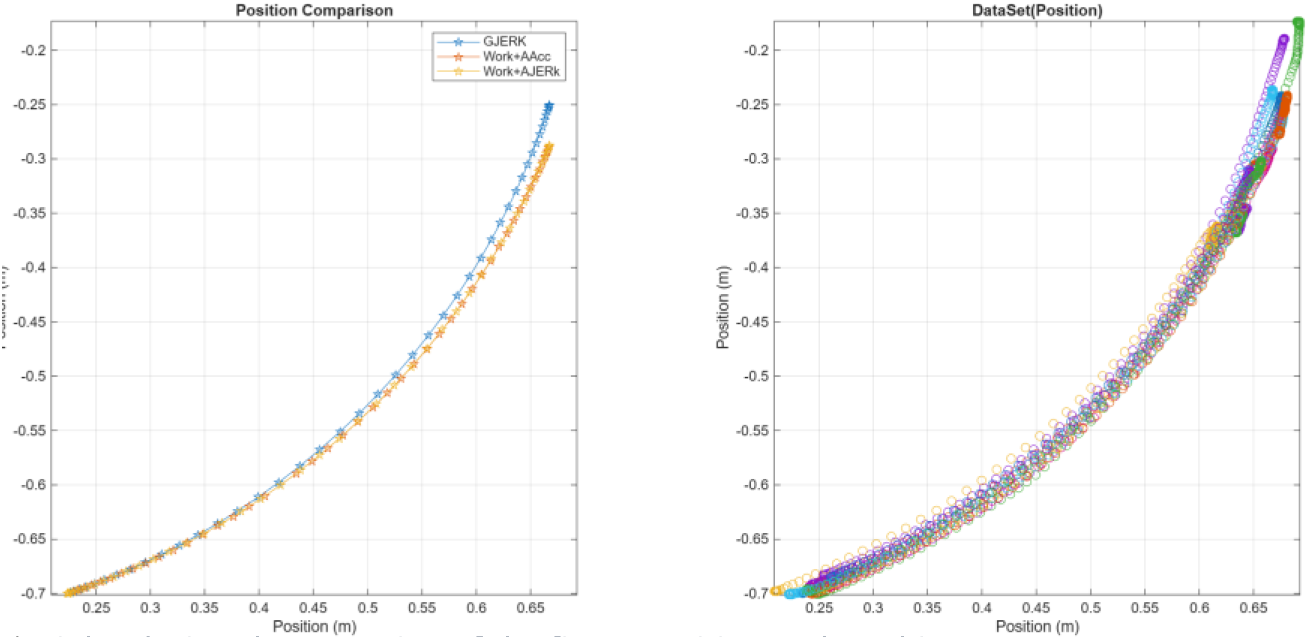
The following figure illustrates the Cost functions trajectories on the left and the actual experimental data trajectories of real human arm movement on the right, during the execution of the first transition P4 by subject S2.

**Figure 24:**
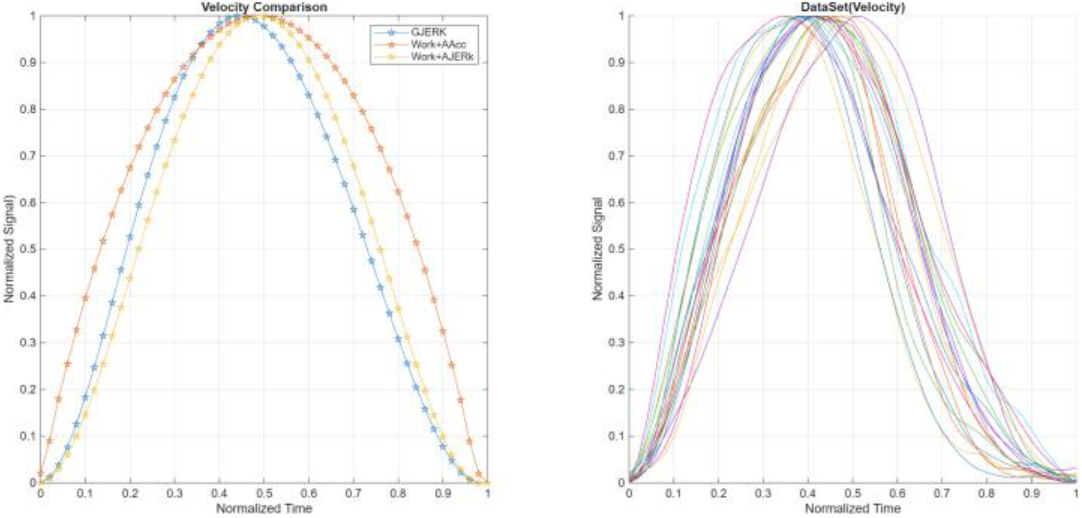
The following figure illustrates the velocity profiles for the Cost functions on the left and the actual velocities of human arm movement on the right, during the execution of transition P4 by subject S2.

**Figure 25:**
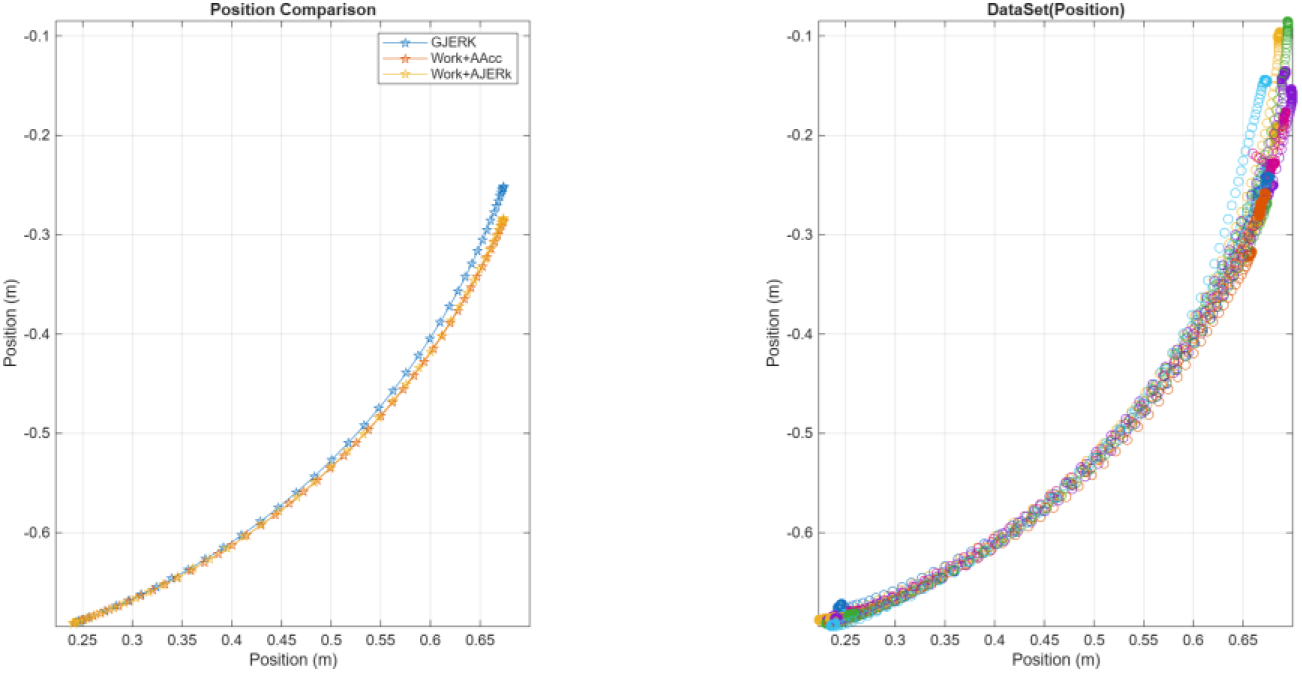
The following figure illustrates the Cost functions trajectories on the left and the actual experimental data trajectories of real human arm movement on the right, during the execution of the first transition P4 by subject S3

**Figure 26:**
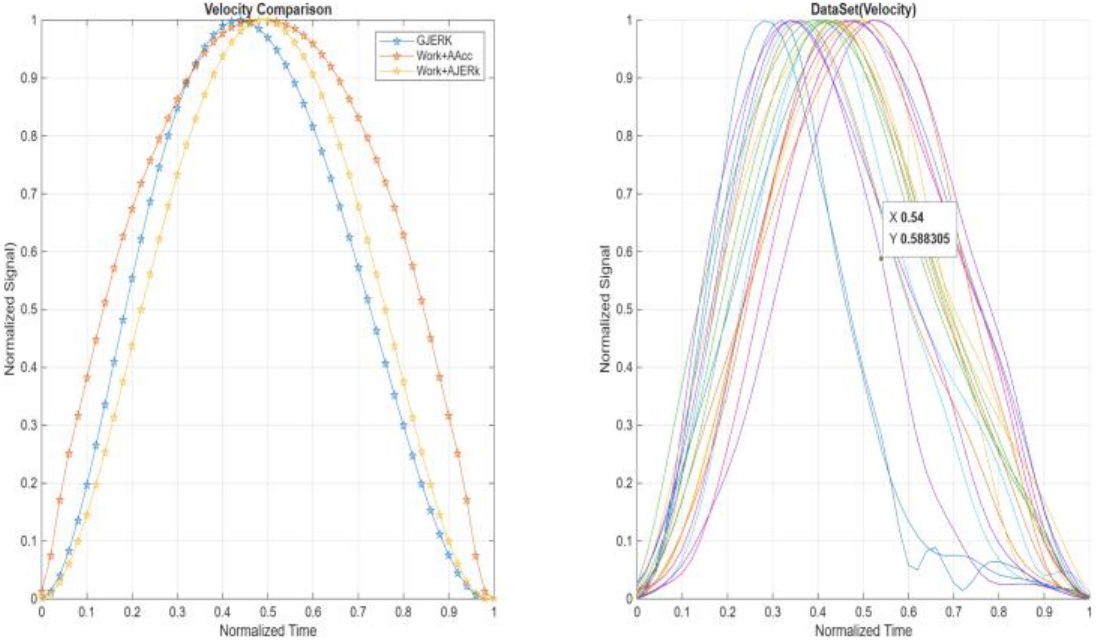
The following figure illustrates the velocity profiles for the Cost functions on the left and the actual velocities of human arm movement on the right, during the execution of transition P4 by subject S3.

**Figure 27:**
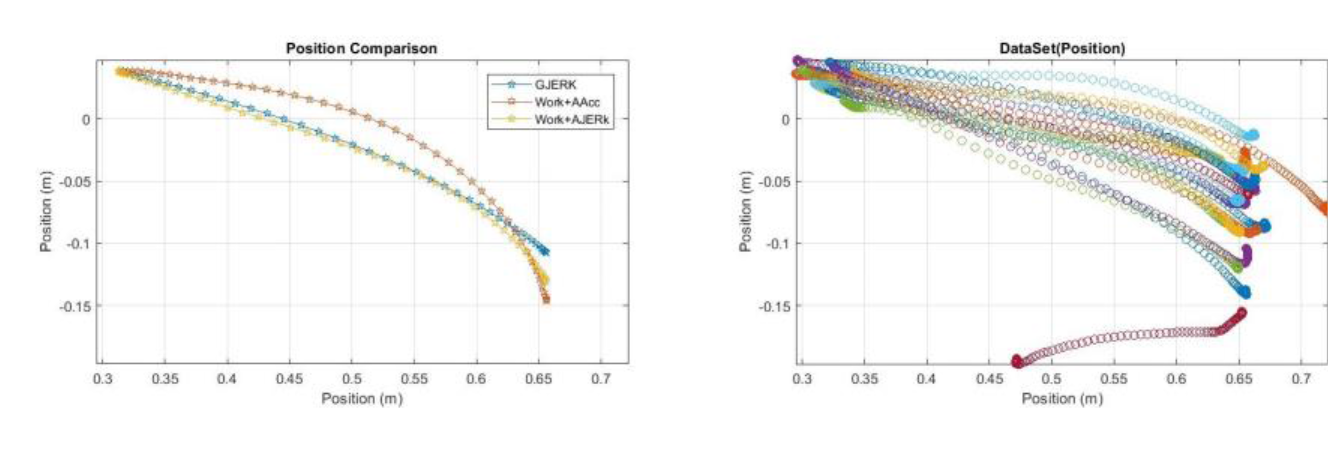
The following figure illustrates the Cost functions trajectories on the left and the actual experimental data trajectories of real human arm movement on the right, during the execution of the first transition P5 by subject S1

**Figure 28:**
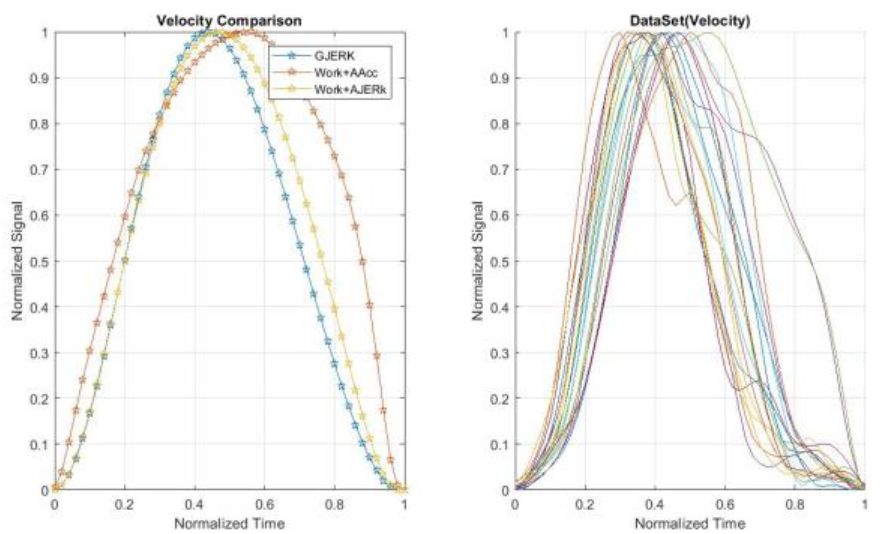
The following figure illustrates the velocity profiles for the Cost functions on the left and the actual velocities of human arm movement on the right, during the execution of transition P5 by subject S1.

**Figure 29:**
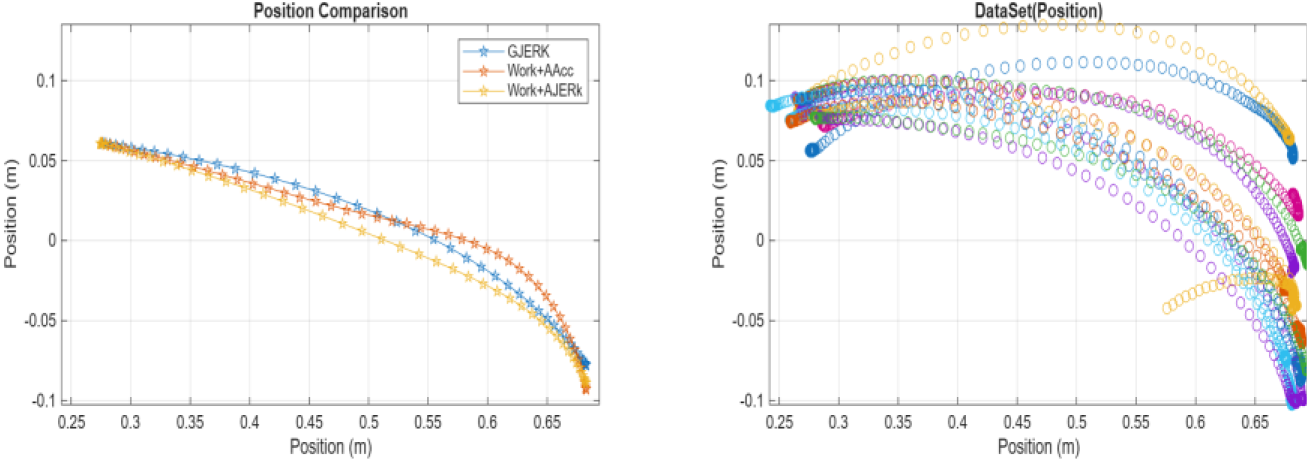
The following figure illustrates the Cost functions trajectories on the left and the actual experimental data trajectories of real human arm movement on the right, during the execution of the first transition P5 by subject S2

**Figure 30:**
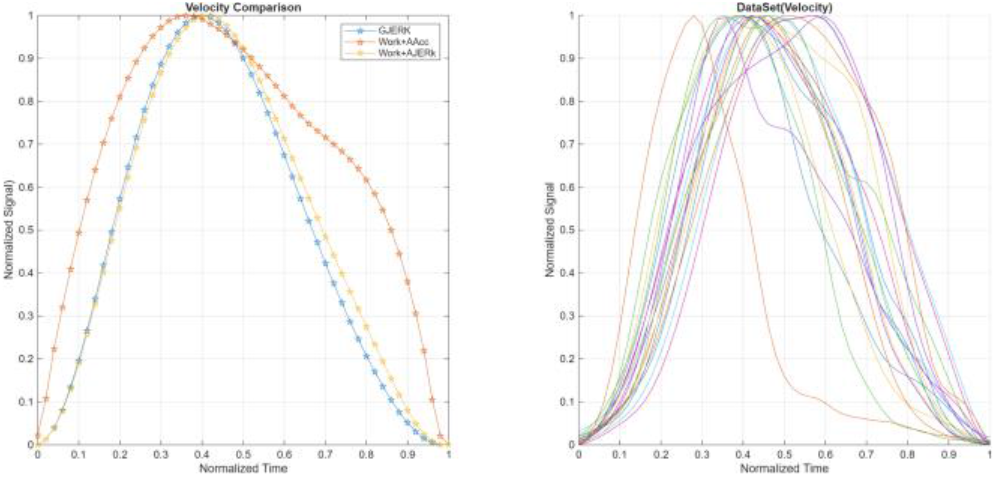
The following figure illustrates the velocity profiles for the Cost functions on the left and the actual velocities of human arm movement on the right, during the execution of transition P5 by subject S2.

**Figure 31:**
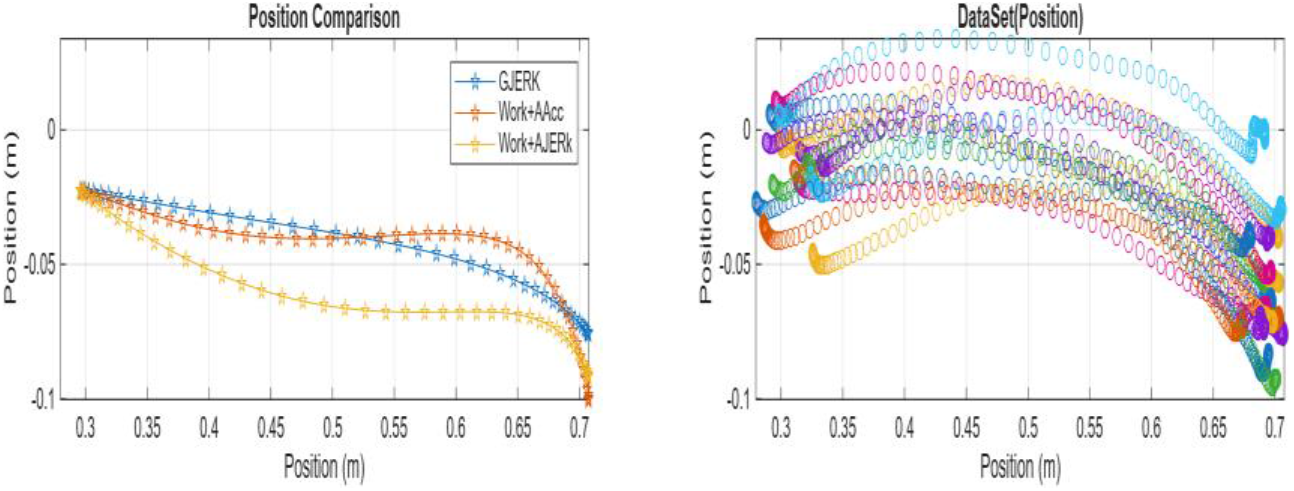
The following figure illustrates the Cost functions trajectories on the left and the actual experimental data trajectories of real human arm movement on the right, during the execution of the first transition P5 by subject S3

**Figure 32:**
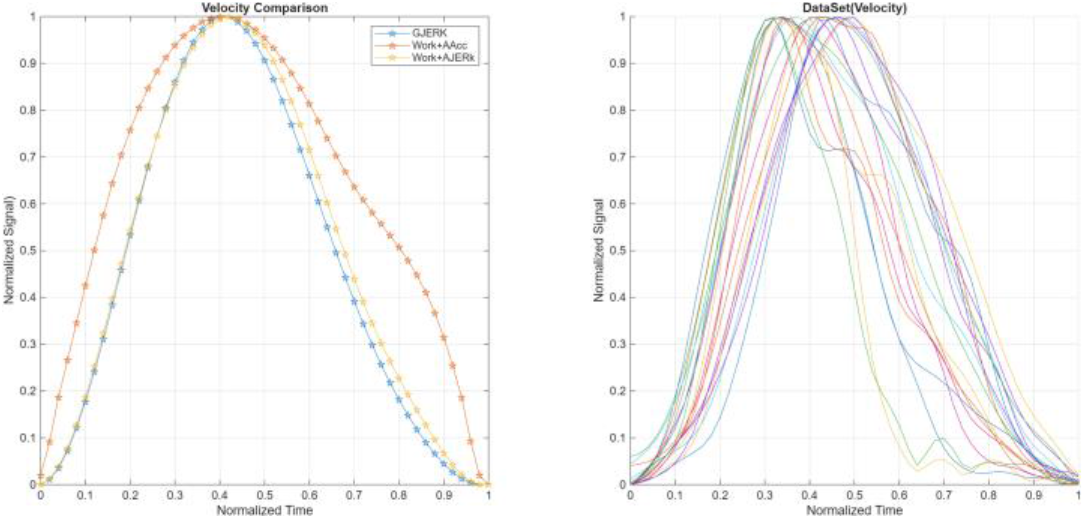
The following figure illustrates the velocity profiles for the Cost functions on the left and the actual velocities of human arm movement on the right, during the execution of transition P5 by subject S3.

Taken together, the expanded set of visualizations and numerical findings provides compelling evidence supporting the validity and reliability of our proposed model. The consistency observed between the quantitative indicators and the detailed trajectory plots across all five movements reinforces the conclusion that the GJERK cost function is not only statistically superior but also dynamically accurate in reproducing the natural behavior of human arm motion. The close alignment between the predicted paths and the experimentally recorded trajectories—both in terms of spatial shape and temporal coordination—demonstrates that our formulation captures the essential biomechanical principles governing voluntary upper-limb movement. Consequently, these integrated results confirm that the GJERK model achieves a high level of precision in describing human arm kinematics, offering a robust and physiologically plausible representation of real motor performance.

Beyond the numerical superiority and the visual agreement demonstrated across all five movements, the collective evidence establishes a coherent and compelling narrative regarding the reliability of the GJERK formulation. The model’s ability to consistently reproduce both the geometric structure and the temporal dynamics of human arm motion—despite the substantial variability inherent in biological trajectories—confirms that its underlying gravity-weighted jerk modulation captures the essential principles governing natural motor behavior. This convergence between theory, quantitative performance, and empirical visualization indicates that the GJERK framework does not merely approximate human movement, but rather provides a physiologically grounded representation of how the neuromuscular system organizes motion under gravitational influence. Consequently, the results affirm that the proposed cost function achieves a high degree of descriptive accuracy, offering a robust and biologically plausible model for characterizing human arm kinematics across diverse movement conditions.

## Discussion

In the vertical plane, the proposed GJERK function demonstrated clear superiority in generating asymmetric linear velocity profiles, a defining characteristic of human motion under gravitational influence. Its gravity-dependent weighting allows the model to redistribute jerk across the joints in a manner that accurately reflects the biomechanical demands of vertical movements, enabling it to reproduce the experimentally observed temporal asymmetry with high fidelity.

In the horizontal plane, however, gravitational torque is completely absent. As a result, the weighting term inside the GJERK formulation collapses to a constant, causing the cost function to effectively reduce to a jerk-only model. This transformation naturally produces symmetric velocity profiles, which aligns precisely with the known kinematic properties of horizontal human movements, where no gravitational bias exists to induce temporal asymmetry. Thus, the GJERK function behaves in a biomechanically consistent manner across planes: it becomes purely jerk-based when gravity has no effect, and regains its dynamic, gravity-modulated structure when vertical motion is involved.

Beyond its biomechanical consistency, the proposed **GJERK** function represents a significant conceptual advancement in the modeling and control of robotic systems. By embedding gravity directly into the weighting structure of the cost function, the formulation provides a principled way to exploit gravitational torque as an assisting force rather than treating it as an external disturbance or constraint. This perspective opens new possibilities in trajectory planning for robots operating in gravity-dominated environments, where the natural gravitational field can be leveraged to reduce actuation effort, smooth motion, and enhance energy efficiency. In this sense, the GJERK formulation offers a powerful framework for designing motion strategies that align with the intrinsic dynamics of the system, enabling robots to benefit from gravity in the same way the human neuromuscular system does—using it as a supportive component rather than an obstacle to be compensated for.

This perspective represents a significant conceptual advancement in the modeling and control of robotic systems. By embedding gravity directly into the cost structure, the GJERK formulation provides a principled way to exploit gravitational torque as an assisting force rather than treating it as a constraint to be overcome. Such an approach opens new possibilities in trajectory planning for robots operating in gravity-dominated environments, where the natural gravitational field can be leveraged to reduce actuation effort, smooth motion, and enhance energy efficiency. In this sense, the GJERK model offers a powerful framework for designing motion strategies that align with the intrinsic dynamics of the system, enabling robots to benefit from gravity in the same way the human neuromuscular system does—using it as a supportive component rather than an obstacle

Moreover, this gravity-aware formulation becomes even more impactful in robotic systems with redundant degrees of freedom. Redundant robots possess more joints than are strictly required to accomplish a given end-effector task, creating a high-dimensional solution space in which internal joint motions can be optimized independently of the primary task objective. By incorporating gravitational torque directly into the optimization landscape, the GJERK model allows these robots to select joint trajectories that make optimal use of gravitational assistance. Instead of counteracting gravity through active compensation, the robot can adopt joint configurations that naturally align with the direction of gravitational support, thereby reducing torque demands, minimizing energy expenditure, and generating smoother, more human-like motion.

This capability fundamentally transforms redundancy resolution from a purely kinematic problem into a dynamically meaningful optimization process. The robot can exploit its extra degrees of freedom to channel gravitational forces in beneficial ways—much like the human motor system does when coordinating multi-joint movements under varying gravitational loads. As a result, the GJERK formulation provides a unified and biologically inspired framework for motion planning in redundant robots, enabling them to harness gravity as a functional resource rather than a limiting factor.

## Acknowledgments

I would like to express my sincere thanks to Professor Bastien Berret, who provided me with the practical data.

## Code and Data Availability

https://doi.org/10.5281/zenodo.21967952

